# Predicting flowering time using integrated morphophysiological and genomic data with machine learning models

**DOI:** 10.1101/2025.08.11.669697

**Authors:** Mehdi Babaei, Hossein Nemati, Hossein Arouiee, Davoud Torkamaneh

**Affiliations:** Department of Horticultural Sciences, Ferdowsi University of Mashhad, Azadi Square, Mashhad, 9177948974, Razavi Khorasan, Iran; Département de Phytologie, Université Laval, Rue de l’Université, Québec City, G1V 0A6, Québec, Canada; Institut de Biologie Intégrative et des Systèmes (IBIS), Université Laval, Rue de l’Université, Québec City, G1V 0A6, Québec, Canada; Centre de recherche et d’innovation sur les végétaux (CRIV), Université Laval, Rue de l’Agriculture Québec City, G1V 0A6, Québec, Canada; Institute Intelligence and Data (IID), Université Laval, Rue de l’Agriculture Québec City, G1V 0A6, Québec, Canada

**Keywords:** Cannabis, Marker-trait Associations, Machine learning, Classification, Flowering Time, Landraces

## Abstract

Indigenous *Cannabis Sativa* populations have adapted to diverse environments, resulting in genetic and phenotypic diversity. Understanding the mechanisms underlying flowering time variation is crucial for optimizing cultivation and breeding. This study employed a novel approach combining temporal phenotypic analysis, genomic data, and machine learning (ML) to identify key features associated with early, medium, and late flowering in cannabis landraces. We collected weekly data on six morphophysiological traits—stem diameter, height, growth rate, node number, internode length, and SPAD chlorophyll index—from 25 cannabis landrace populations 13 weeks for female plants and 11 weeks for male plants. Additionally, 145 accessions were genotyped using high-density genotyping-by-sequencing, resulting in 233,624 high-quality single nucleotide polymorphisms (SNPs). A comprehensive ML framework integrating mutual information (MI), recursive feature elimination (RFE), random forest (RF), and support vector machine (SVM), was used to investigate 234,002 features, encompassing SNPs, morphophysiological traits, and environmental factors. This approach identified 53 key features—22 genetic variants and 31 morphophysiological traits—that effectively distinguish between early, medium, and late flowering types with an accuracy of 96.6%. The identified SNPs were distributed across multiple chromosomes, including chromosomes 08, 09, and X. Notably, key loci like *AutoFlower3* (*CsFT3*) (on chromosome 08) and *CircadianFloweringLocus1* (*CsCFL1*) (on chromosome 09) were identified, with several SNPs located within or near annotated genes. These findings contribute significantly to the understanding of cannabis chronobiology and support the development of “smart crop” strategies by providing valuable markers for early selection and targeted breeding programs aimed at optimizing flowering time under diverse conditions.

**Key Message:** A data-driven machine learning strategy combining genomic and dynamic phenotypic traits enables accurate classification of flowering time in diverse *Cannabis* landraces.

## 1. Introduction

Cannabis (*Cannabis sativa* L.), commonly known as hemp, marijuana, marihuana, Indian hemp, and industrial hemp, is among the earliest domesticated and cultivated plants in human history (Crocq 2020; Babaei et al. 2022). Its long history is closely intertwined with human evolving social, cultural, and economic developments, marked by shifts over time (Barcaccia et al. 2020). Cannabis has been an integral part of human life since its domestication, although studies on its use have substantially decreased since the mid-20^th^ century, when it became illegal in most Western countries (Warf 2014). As a result, cannabis has not completely benefited from modern scientific advances and advanced breeding techniques, resulting in a massive knowledge gap that must be rectified. With shifting legislation and increasing legalization in countries such as the Uruguay Canada, United States, the Netherlands, Germany, Thailand, and Morocco the cannabis market continues to expand steadily. The global cannabis market is forecasted to reach a value of US$75.09 billion by 2029, driven by ongoing developments in innovation and product diversification (Statista 2024).

Cannabis is an annual flowering member of the order Urticales and the family Cannabaceae, which includes another well-known genus *Humulus* (*H. lupulus*, *H. yunnanensis*, and *H. japonicus*) (Ii 2003; Shephard et al. 2004; Ahmed et al. 2008; Kovalchuk et al. 2020; Schilling et al. 2020). The taxonomy of the *Cannabis* genus has been a subject of ongoing debate. It is unclear whether it is monospecific (*C. sativa*) or poly-specific (*C. sativa*, *C. indica*, and *C. ruderalis*) (Small and Beckstead 1973; Anwar et al. 2006; Flores-Sanchez and Verpoorte 2008; McPartland and Guy 2017; McPartland 2018; Schwabe and McGlaughlin 2019). However, the diversity of modern cannabis cultivars often extends beyond legacy taxonomic definitions (*sativa*, *indica*, and *ruderalis*) due to years of undocumented hybridization. Cannabis cultivars and *forma* are defined within distinct clades, including legal status, phytochemical profile (chemotype), ecological adaptation (ecotype), and biological characteristics (biotype) (Babaei and Ajdanian 2020; Lapierre et al. 2023; Babaei et al. 2025). One key distinction is based on delta-9-tetrahydrocannabinol (THC) concentration, which separates hemp with low THC content (< 0.3%) in dry-weight flower buds from drug-type cannabis (THC: CBD ratio > 1 THC-predominant with CBD stands for cannabidiol) (De Meijer and Hammond 2005; Sawler et al. 2015; Lapierre et al. 2023).

Cannabis is predominantly dioecious, with male (XY) and female (XX) plants, a diploid genome of 2n = 20, though monoecious forms and hermaphroditism can occur under specific conditions (Carpentier et al. 2012; Braich et al. 2020). Recent studies have revealed that ethylene biosynthesis genes influence sexual plasticity—the ability to alter phenotypic sex without chromosomal changes—in cannabis, particularly under environmental stress or chemical treatments such as silver thiosulfate (STS), which inhibit ethylene signaling (Monthony et al. 2024).

Cannabis is generally a short-day plant, with flowering triggered by photoperiods of 12 to 15 hours (Lisson et al. 2000; Amaducci et al. 2008; Zhang et al. 2021). However, photoperiod-insensitive (day-neutral or auto-flower) cultivars, often associated with *C. sativa* var *ruderalis*, flower regardless of day length, likely as an adaptation to high-latitude conditions (McPartland 2018). Flowering times vary significantly among cannabis cultivars due to strong genetic control. In plants, key regulators influencing flowering behavior include genes, transcription factors, small RNAs, and epigenetic marks (Kumari et al. 2022; Shi et al. 2022a, 2023; Ibrar et al. 2023; Naik et al. 2025). Overall, plants integrate environmental signals like photoperiod and temperature through these genetic networks to fine-tune flowering. (Amaducci et al. 2005; Salentijn et al. 2015; Cao et al. 2021; Babaei et al. 2022, 2024; Osnato et al. 2022). Flowering time is a critical factor in hemp breeding, as peak fiber quality is reached shortly after flowering (Amaducci et al. 2005, 2008; Salentijn et al. 2015; Petit et al. 2020a, b). Flowering time also plays a pivotal role in cannabinoid production, as cannabinoids accumulate rapidly during the early (the first three weeks) flowering stages (Stack et al. 2021). Delaying flowering can increase cannabinoid yields in some genotypes, but this often comes with a trade-off between floral biomass and cannabinoid concentration (Steel et al. 2023). However, the strong genetic control over this trait has enabled selection of early-flowering (auto-flowering) cultivars for short season seed production, and late-flowering types for fiber production in high-latitude areas (Salentijn et al. 2019). While breeding efforts have focused on selecting lines with flowering times appropriate for the intended use, optimizing these processes requires a deeper understanding of the genetic mechanisms underlying flowering (Dowling et al. 2021, 2024).

The key genetic factors influencing flowering time in cannabis have recently been identified. Toth et al. (2022) identified two significant loci: *Autoflower1*, associated with photoperiod insensitivity, and *Early1*, which promotes early flowering. Similarly, Dowling et al. (2024) linked the photoperiod insensitivity of the “FINOLA” hemp cultivar to a duplication of the *FLOWERING LOCUS T* (*FT1*) gene, termed *Autoflower2*. Petit et al. (2020) further identified multiple quantitative trait loci (QTL) associated with flowering time, suggesting the involvement of pathways regulating photoperiod sensing, circadian rhythms, and hormone signaling. These insights provide a foundation for tailoring flowering traits to specific breeding objectives. However, the integration of genetic insights with phenotypic and environmental data remains underexplored, especially in diverse landrace populations.

Recent studies have advanced cannabis research by integrating analytical and modeling techniques to enhance the understanding of flowering time, growth patterns, and chemitypic classification. Hyperspectral imaging (HSI) combined with multivariate data analysis achieved a classification accuracy of 94.7% for identifying different cannabis chemotypes (San Nicolas et al. 2024). Similarly, polynomial models to analyze nonhomologous landmarks, enabling accurate predictions of accession identity, leaflet number, and node position across nine cannabis accessions (Balant et al. 2024). Research on weekly growth trends and chemical changes has further refined predictive modeling approaches. A study with 121 genotypes under controlled conditions established correlations between early-stage growth traits—such as plant height and stem diameter—and dry biomass weight, allowing multivariate regression models to predict biomass based on vegetative growth rates (Naim-Feil et al. 2021). In parallel, a field study on 30 hemp cultivars monitored weekly changes in height, flowering time and cannabinoid accumulation (CBD, THC, and cannabichromene (CBC)). By using regression models for growth rates, polynomial models for cannabinoid accumulation, and linear models for biomass estimation, researchers optimized cultivation strategies to enhance yield and chemical composition (Stack et al. 2021). These advancements highlight the role of innovative modeling and analytical techniques in improving cannabis research, particularly in predicting flowering time and optimizing biomass production.

This study focuses on understanding the diversity in flowering times among cannabis landrace populations through a comprehensive analysis of phenotypic, genomic, and environmental data. Using 25 Iranian landrace populations previously classified as early, medium, and late flowering (Babaei et al. 2024), we tracked six morphophysiological traits weekly across 13 weeks for female and 11 weeks for male plants. Expanding this investigation to 145 native accessions, we utilized genomic sequencing and machine learning (ML) to analyze a total of 234,002 features, identifying key genetic variants associated with flowering time variation. The findings from this study not only provide a detailed understanding of flowering-related traits but also offer practical tools for early selection and classification of flowering types, facilitating breeding programs and optimizing cultivation strategies for cannabis.

## 2. Materials and methods

### 2.1 Plant materials

Cannabis seed samples from 25 native populations (Table S1) of Iran were sourced from local markets and native farmers in various regions, ensuring genetic diversity and regional adaptation. The regions from which the seeds were collected encompassed five defined climatic zones based the Köppen-Geiger climate classification method, with geographic latitudes between 25° and 40° North and Longitudes 45° and 65° East. Prior to cultivation, preliminary germination tests were conducted to evaluate key parameters (Babaei et al. 2024).

### 2.2 Experimental design and growing conditions

Twenty seeds from each collected population were planted (Babaei et al. 2024). Thirty days after sowing (DAS), the seedlings were transferred to growth bags with a mixture consisting of garden soil, leaf mold, sand, and perlite in a 3:1:1:1 proportion. Based on the study by Amaducci et al. (2008), the planting was done 60 days before the photoperiod switch-off in Mashhad, Khorasan, Iran when the days start to shorten (the day length at sowing was 13 hours and 17 minutes) to ensure that the plants would have an adequate vegetative period before entering the reproductive phase. As shown in Fig. 1, on the 60^th^ DAS the maximum day length (14 hours and 29 minutes) was considered the start of the reproductive phase, with the period before it being regarded as the vegetative phase.

**Fig. 1.**
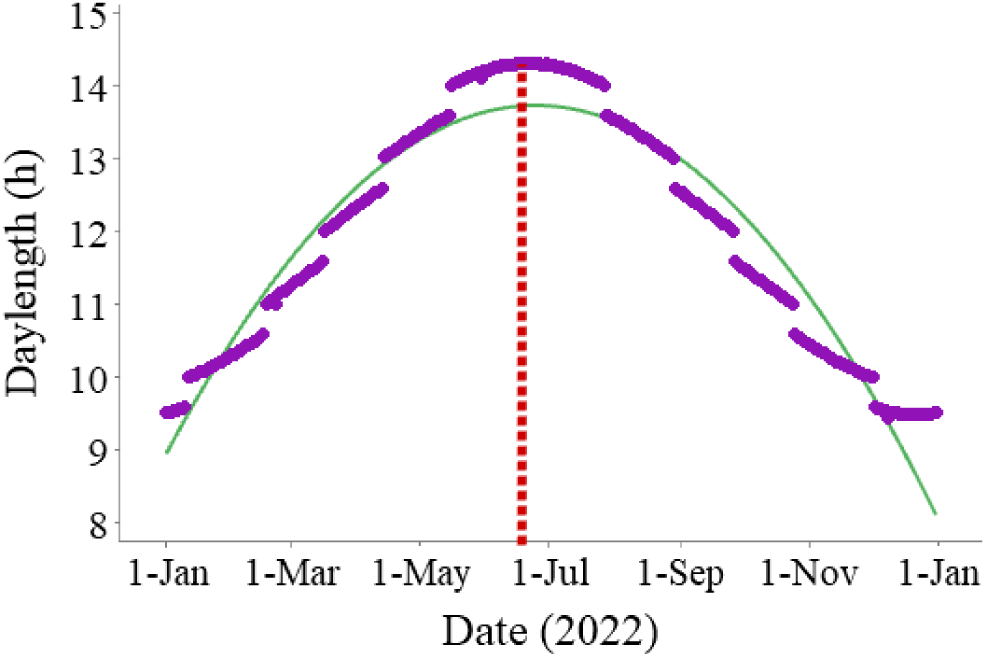
Day-length variation throughout the year (2022) in Khorasan, Iran. The red line indicates the photoperiod Switch-Off, marking the start of day-length shortening.

The plants were arranged in a randomized complete block design (RCBD) with three blocks and five observations, totaling 375 plants, in the greenhouse complex of Ferdowsi University of Mashhad, Iran (36^°^16’N and 59^°^36’E with an altitude of 985 m). According to Spatial Analysis (SA), each unit, measuring 60 square meters, was divided into 25 rows and 15 columns, so that each block included 5 columns. The aisles within each unit were arranged so that 16 plants were placed per square meter. At the beginning of the reproductive stage (appearance of solitary flowers), male and female plants were separated and transferred to a unit with similar conditions, reducing the number of plants per square meter by half. The growing conditions and all cultural practices were previously described in Babaei et al. (2024).

### 2.3 Data collection

#### 2.3.1 Determination of morphophysiological traits

The first data collection was conducted after plants establishment (30 DAS), followed by weekly, individual measurements carried out for 13 weeks on female plants and 11 weeks on male plants. Since the early-flowering populations (IR7385 and IR2845) had a very short growth period, they had a completely different growth pattern. For this reason, female plants were harvested in the 10^th^ week of data collection (93 DAS), while male plants were harvested in the 6^th^ week (63 DAS).

The measured traits in this experiment included Stem Diameter (SD), plant Height (H), Growth Rate (GR), the Number of Nodes (NN) on the main stem, Length of Internode in th Middle Third of the Main Stem (LI-MT), and SPAD-based chlorophyll (SPAD). The SD at 5 cm above soil surface was measured using a caliper with an accuracy of 0.01 mm. The H and LI-MT were measured using a tape with an accuracy of 0.01 m. In addition, the NN on the main stem was counted. The chlorophyll index was measured using a SPAD-502 meter (Konica, Minolta, Tokyo), with SPAD readings taken on three leaves (top, middle, and bottom) of each plant, and the average was calculated for each individual. Finally, the GR, expressed in centimeters per day (cm/day), was calculated through the following equation (eq. 1) (Stack et al. 2021):

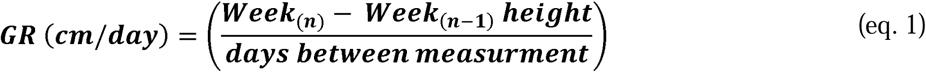

#### 2.3.2 Genotyping

Sampling was performed from healthy and young leaves of all individuals. Samples were then dried using the Freeze Dry System Alpha 2-4 LD plus for 24 hours. A total of 145 pooled samples, with each sample representing a unique combination of males and females from each population, were ground with metallic beads in a RETSCH MM 400 mixer mill (Fisher Scientific, MA, USA). DNA extraction was carried out using the Qiagen DNeasy^®^ Plant Mini Kit. The extracted DNA quantity was assessed by a Qubit fluorometer with the dsDNA HS assay kit (Thermo Fisher Scientific, MA, USA), and their quality was randomly evaluated using agarose gel electrophoresis. DNA concentrations were adjusted to 10 ng/μl for all samples. Final DNA samples were used to prepare HD-GBS libraries with *BfaI* as described in Torkamaneh et al., (2021) at the Institut de biologie intégrative et des systèmes (IBIS), Université Laval, QC, Canada. Sequencing was conducted on an Illumina NovaSeq 6000 with 150 paired end reads at the Genome Quebec Service and Expertise Center (CESGQ), Montreal, QC, Canada.

#### 2.3.3 SNP calling and filtration

Sequencing data were processed with the Fast-GBS v2.0 using the *C. sativa* cs10 v2 reference genome (GenBank acc. no. *GCA_900626175.2*) (Grassa et al. 2018; Torkamaneh et al. 2020). For variant calling a prerequisite of a minimum of 6 reads to call a single nucleotide polymorphism (SNP) was opted. Raw SNP data were then filtered with VCFtools to remove low-quality SNPs (QUAL <10 and MQ <30) and variants with proportion of missing data exceeding 80% (Danecek et al. 2011). Missing data imputation was performed with BEAGLE 4.1 (Browning and Browning 2016), followed by a second round of filtration, retaining only biallelic variants with heterozygosity less than 50% and a minor allele frequency (MAF) of > 0.05. Additionally, variants residing on unassembled scaffolds were removed (de Ronne et al. 2024).

### 2.4 Machine learning (ML) datasets

We employed the ML framework introduced by Abdelwahab et al. (2022) to identify the most important features for distinguishing the three flowering time clades. This framework is schematically represented in Fig. 2. The populations were categorized into three flowering clades based on the number of DAS to the “Start 10% Flowering Time in Individuals” (10% of bracts formed) (SF10I): early (60-80 days for females, 40-60 days for males), intermediate (80-115 days for females, 60-95 days for males), and late (115-140 days for females, 95-120 days for males) (Babaei et al. 2024).

**Fig. 2.**
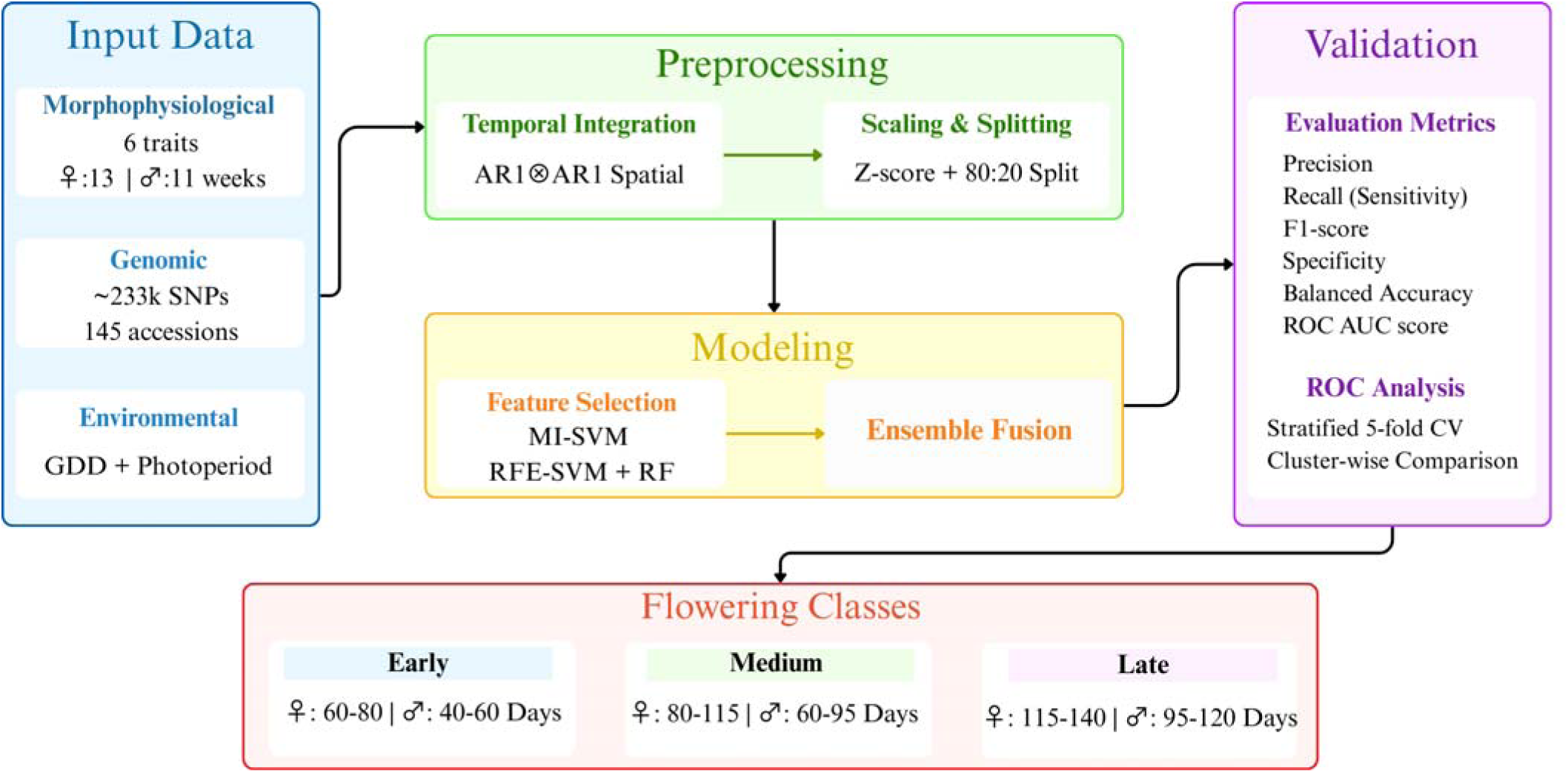
A graphical overview of the proposed model’s framework architecture for feature identification in the classification of various phenological stages.

The dataset used in this framework included data on six key morphophysiological traits (SD, H, GR, NN, LI-MT, and SPAD) collected over 13 weeks for female plants and 11 weeks for male plants, along with catalog of 233,624 high-quality SNPs obtained for 145 accessions: 73 male and 72 female plants. This accession panel comprised 12 early-flowering, 66 medium-flowering, and 67 late-flowering accessions. Environmental data, including the average daily temperature based on growing degree-days (GDD ^°^C) as described in equation (eq. 2) and day length over 150 days, were also used. In total, 234,002 features were incorporated and analyzed within this framework.

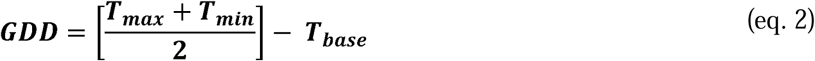

Where *T_max_*is the daily maximum temperature (°C), *T_min_* is daily minimum temperature (°C) and *T_base_* is the base temperature of the leaf emerging, where 1 ^°^C was considered (Cannoy 2015).

#### 2.4.1 Data Preprocessing

The weekly data for six morphophysiological traits were fitted using SA and combined with the SNP data. The features were normalized to have zero mean and unit variance using the following equation (eq. 3). To train and test the models, 80% of the data (116 accessions: 10 early-flowering, 53 medium-flowering, and 53 late-flowering) were used for training, and 20% (29 accessions: 2 early-flowering, 13 medium-flowering, and 14 late-flowering) were used for testing the models (Fig. 2).

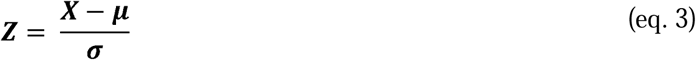

where:

*X* = The single raw data value

*μ* = The mean

*σ* = The standard deviation

#### 2.4.2 Machine learning (ML) algorithms

In this framework, four ML algorithms were employed: Mutual Information (MI), Support Vector Machine (SVM), Recursive Feature Elimination (RFE), and Random Forest (RF) (Kraskov et al. 2004; Jakkula 2006; Chen and Jeong 2007; Belgiu and Drăguţ 2016). The MI examines the relationship between features and classes, as well as the redundancy among features. To address the issue of binning, an estimator with K-neighbors (k=3) was used. In this framework, MI, as a filter-based Feature Selection (FS) technique, initially sorted features based on their importance value. Features with very low or zero MI were placed at the bottom of the list. Consequently, the top 1,000 features were selected and evaluated in SVM (MI-SVM) to determine the optimal feature set. SVM with a linear kernel was employed to evaluate the classification performance of features selected by MI and RFE.

The RFE model, a wrapper-based FS technique, was used to identify the optimal subset of features that can achieve the best and highest performance by creating a subset of features at each stage and retaining the best subset. Features were then ranked based on their removal. The criterion for removal (i.e., SVM with a linear kernel (RFE-SVM)) was applied in the predictive model. In this framework, the algorithm was set to remove 50% of features in each iteration. Specifically, the training data was divided into two equal parts in each iteration, and the features that achieved the highest score in SVM were retained.

The RF model, which is based on the embedded feature selection strategy, was applied in this framework. Unlike the filter-based (MI) and wrapper-based (RFE) methods, feature selection in RF model was part of the model-building process. This model created multiple decision trees through the Classification and Regression Trees (CART) method, combined with the bagging technique (Prasad et al. 2006). Each decision tree resulted in a specific prediction, and in this framework, up to 1,000 trees were used to identify the best subset of features (Abdelwahab et al. 2022).

#### 2.4.3 Model evaluation metrics

During training and testing, the ML algorithms were evaluated on several factors, including their precision, recall (sensitivity), F1-score, specificity, balanced accuracy, Receiver Operating Characteristics Areas Under the Curve (ROC AUC) score and confusion matrix, which can be seen in equations (4) – (8):

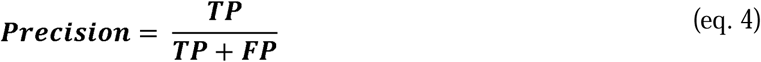

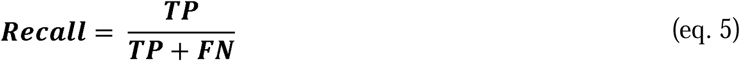

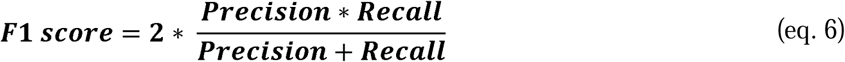

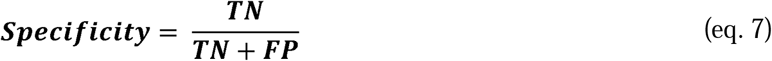

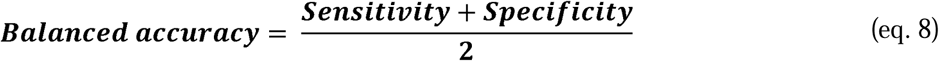

Where:

*TP* (True Positives) = The number of accessions correctly classified as belonging to a specific flowering cluster.

*FP* (False Positives) = The number of accessions incorrectly classified as belonging to a specific flowering cluster.

*TN* (True Negatives) = The number of accessions correctly classified as not belonging to a specific flowering cluster.

*FN* (False Negatives) = The number of accessions incorrectly classified as not belonging to a specific flowering cluster when they belong to that cluster.

More specifically, the ROC AUC metric measures the model’s ability to distinguish between different flowering clusters by evaluating the trade-off between True Positive Rate (TPR) and False Positive Rates (FPR), providing a single, interpretable score (Romagnoni et al. 2019; Khalili et al. 2020; Senan et al. 2021). The confusion matrix offers a detailed breakdown of the model’s predictions, showing how well each accession is classified into the correct flowering cluster and highlighting areas of misclassifications (Alejandrino et al. 2020). Additionally, we used ROC AUC analysis with stratified 5-fold cross-validation to further evaluate each identified marker’s diagnostic potential in the selected feature sets for the SVM classification model, comparing the overall performance across all three clusters and within each cluster. Due to class imbalance in the test set (2 Early, 13 Medium, 14 Late), weighted averages were used for precision, recall, F1-score, and ROC AUC to prioritize majority-class performance. Macro averages were applied for specificity and balanced accuracy to ensure equal representation of all classes. This hybrid approach balances practical relevance (weighted) and fairness (macro) in model evaluation.

### 2.5 Candidate region identification and gene annotation

To identify genomic regions associated with flowering time variation, linkage disequilibrium (LD) and haplotype block (HB) analysis were conducted using PLINK v1.90b5.3 (Purcell et al. 2007). Pairwise LD (*r²*) values were calculated for all biallelic SNPs across each chromosome using the parameters *‘– r2 –ld-window-r2 0’*, enabling full-range LD estimation. HBs were inferred with the *‘–blocks no-pheno-req –ld-window-kb 999’* setting, which defines LD blocks without requiring phenotypic data and allows wide genomic windows for block detection. The results were further processed for visualization using Haploview v4.1, with input formats converted using PLINK’s *‘–recode HV’* function (Barrett et al. 2005). Each HB was delimited by its most upstream (5’) and downstream (3’) SNP positions. Genes located within these intervals were considered putative candidates for regulating flowering behavior.

For gene annotation, functional interpretation of candidate regions was performed using SnpEff v5.2e, based on the *Cannabis sativa* reference genome (cs10 v2; GenBank accession: *GCA_900626175.2*) (Cingolani et al. 2012; Grassa et al. 2018). Gene ontology (GO) terms were extracted from the NCBI *Cannabis sativa* Annotation Release 100 to characterize the potential biological roles of the candidate genes.

### 2.6 Statistical analysis and multiple regression model

The data were first evaluated using the Outlier Grubbs approach to detect and remove any outliers (Adikaram et al. 2015). Subsequently, Minitab^®^ Statistical Software 21 was employed to conduct skewness and kurtosis analyses, assessing the normality of the data and residual errors based on the distribution type (Cain et al. 2017; Minitab 2021). Missing data were imputed using the CART method implemented in the *AllInOne* package in R software version 4.3.1 (Najafabadi et al., 2023; R Team, 2020). SA (AR1⊗AR1) was applied to account for potential spatial effects and minimize environmental impact (i.e., 15 columns and 25 rows). The data fitting approach considered population as a fixed factor and block as a random factor. The model’s performance was evaluated using four primary metrics: Root Mean Square Error (RMSE), Mean Squared Error (MSE), Normalized Root Mean Squared Error (NRMSE), and Coefficient of Variation (CV). Further details and results for these metrics can be found in Table S2. The equations for these metrics are provided below (9) – (12):

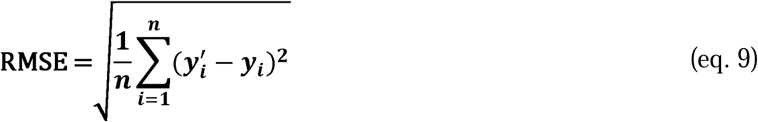

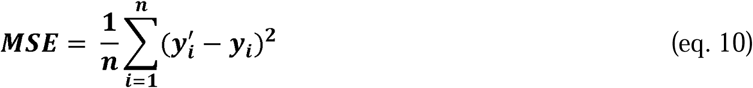

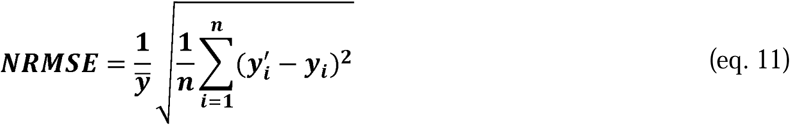

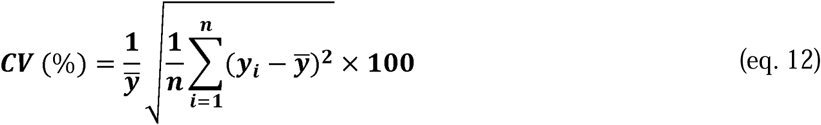

where:

*n* = The number of observations

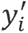 = The predicted value in observation *i*

*y_i_* = The actual value in observation i

*y̶* = The mean of the data values

Broad-sense heritability (H^2^) was determined by the *AllInOne* package (Najafabadi et al. 2023) using following equation (13):

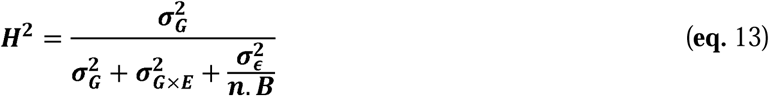

where:

*σ_G_* = The genetic variance

*σ_G×E_ =* The variance due to genotype-by-environment interactions

*σ_ε_ =* The residual variance

*n. B =* The number of blocks

Additionally, the Pearson method and the package *corrplot* (Wei et al. 2017) in R were employed to investigate the correlation between traits. The ML framework was executed using Python 3.12.4, while regression models were developed in MATLAB software R2024a to demonstrate patterns in six morphophysiological traits (the MathWorks Inc., Natick, MA, USA).

## 3. Results

### 3.1 Weekly trends in morphophysiological traits in male and female cannabis plants

We examined the weekly trends of six morphophysiological traits (SD, H, GR, NN, LI-MT, and SPAD index). Data were collected over 13 weeks for females and 11 weeks for males, starting 30 days after sowing (DAS) and ending one month before harvest, except for the early-flowering populations, where data was collected for 9 weeks for females and 5 weeks for males. Based on the averages across 25 native cannabis populations from four different perspectives: *i.* the analytical temporal framework, focusing on population variance (Figs. 3 and 4); *ii.* trait distribution, analyzing changes between vegetative (VS) and reproductive stages (RS) (Fig. 5); *iii.* analysis based on averages, exploring growth pattern differences between female and male plants (Fig. 6); *iv.* broad-sense heritability (H^2^), examining the overall genetic contribution to trait variation (Table 1). Additionally, the heatmap depicting trait correlations is provided in the supplementary materials (Figs. S1 and S2).

**Fig. 3.**
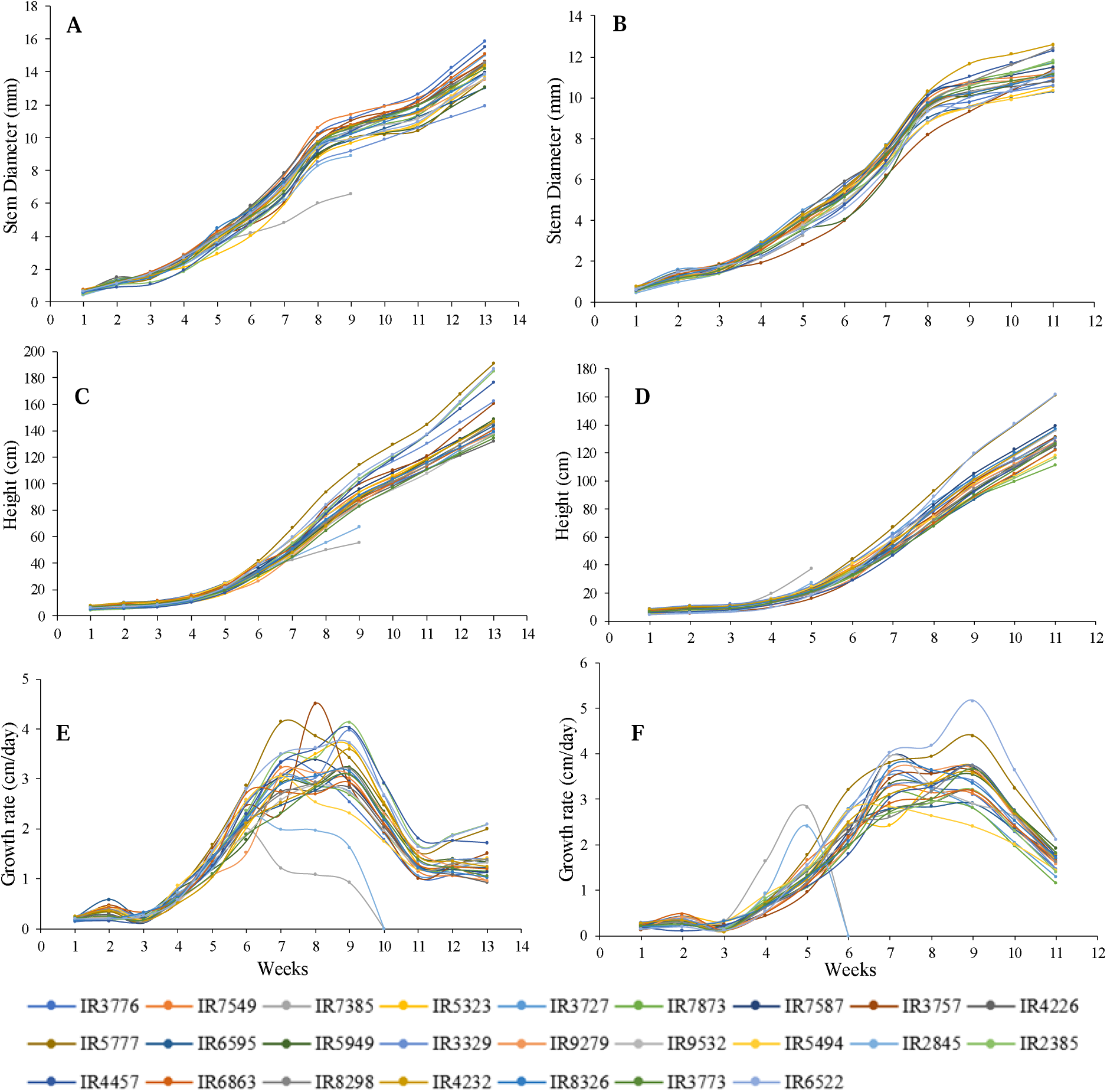
Trends in key morphophysiological traits in 25 native cannabis populations based on Stem Diameter (SD) in female plants (**A**) and male plants (**B**), Height (H) in female plants (**C**) and male plants (**D**), Growth Rate (GR) in female plants (**E**) and male plants (**F**).

**Fig. 4.**
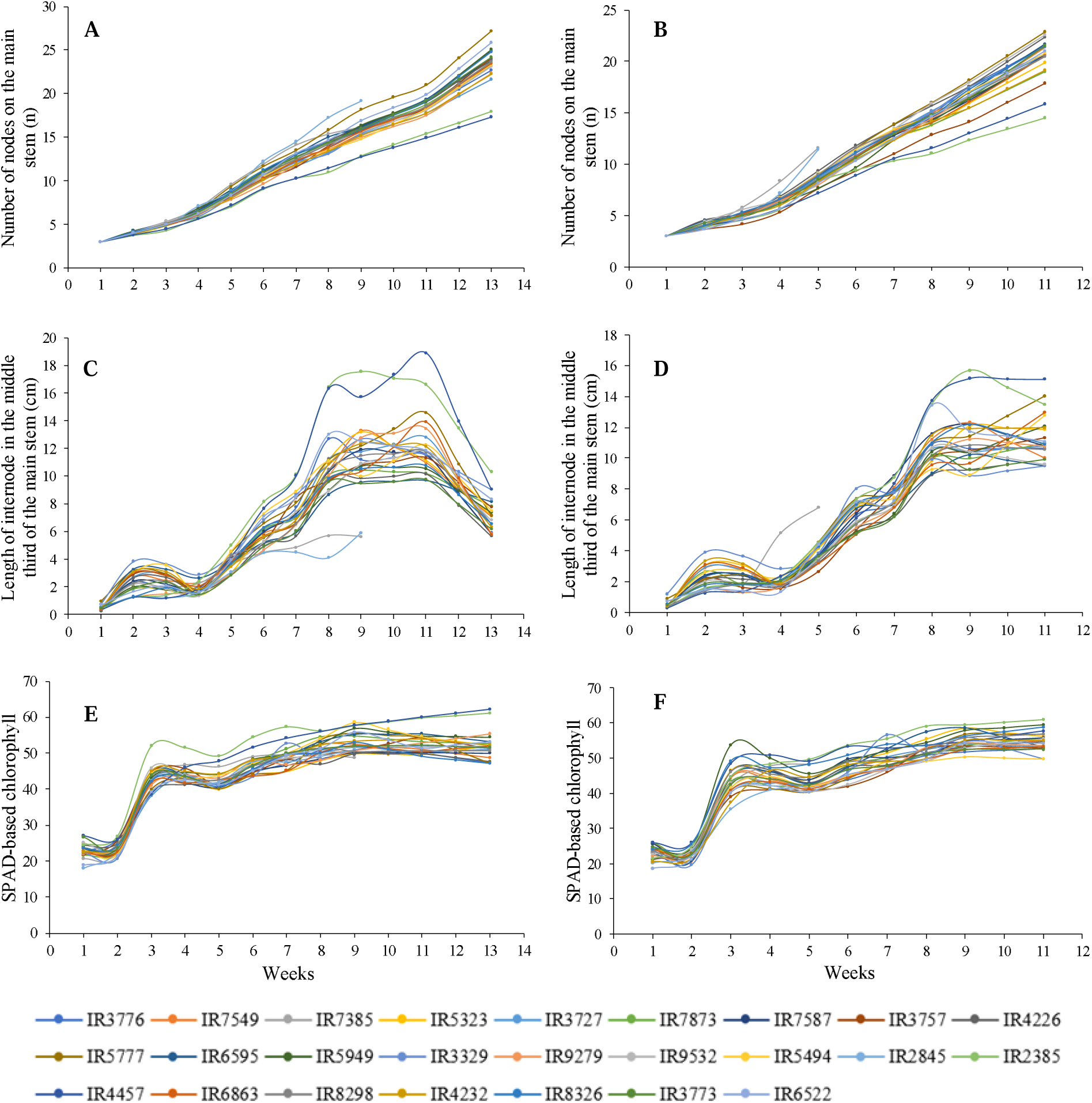
Trends in key morphophysiological traits in 25 native cannabis populations based on Number of Nodes on the Main Stem (NN) in female plants (**A**) and male plants (**B**), Length of Internode in the Middle Third of the Main Stem (LI-MT) in female plants (**C**) and male plants (**D**), SPAD-based Chlorophyll (SPAD) in female plants (**E**) and male plants (**F**).

**Fig. 5.**
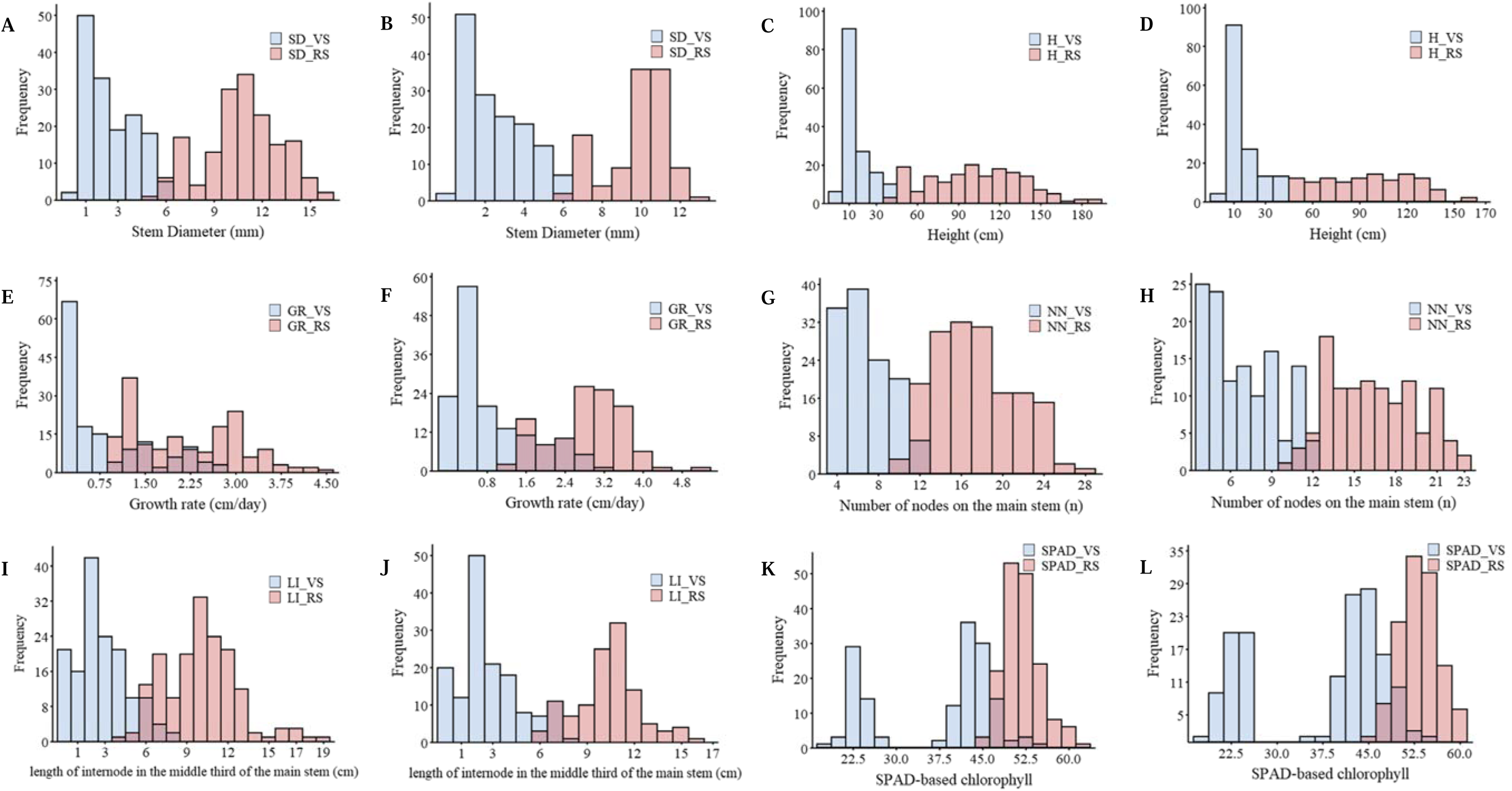
The frequency distributions and range of six key morphophysiological traits based on the averages of 25 native cannabis populations for male and female plants, emphasizing the comparison between the vegetative (VS) and reproductive stages (RS), distinctly separated by the sunlight switch-off. Stem Diameter (SD) in female plants (**A**) and male plants (**B**), Height (H) in female plants (**C**) and male plants (**D**), Growth Rate (GR) in female plants (**E**) and male plants (**F**), Number of Nodes on the Main Stem (NN) in female plants (**G**) and male plants (**H**), Length of Internode in the Middle Third of the Main Stem (LI-MT) in female plants (**I**) and male plants (**J**), SPAD-based Chlorophyll (SPAD) in female plants (**K**) and male plants (**L**).

**Fig. 6.**
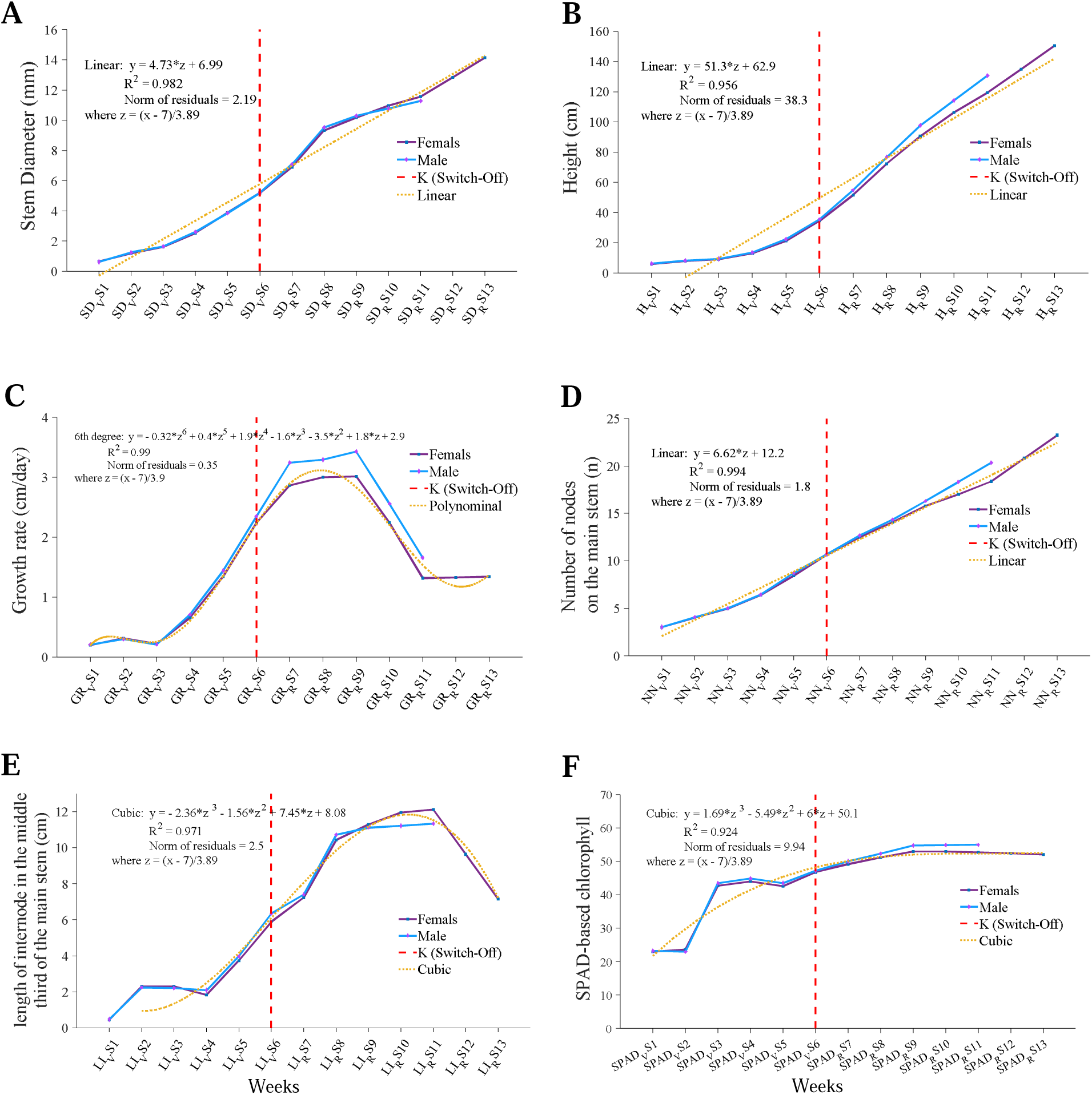
Comparative trends in six key morphophysiological traits based on the averages of 25 native cannabis populations for male and female plants. Traits include (**A**) Stem Diameter (SD), (**B**) Height (H), (**C**) Growth Rate (GR), (**D**) Number of Nodes on the Main Stem (NN), (**E**) Length of Internode in the Middle Third of the Main Stem (LI-MT), and (**F**) SPAD-based Chlorophyll (SPAD). Abbreviations; VS: Vegetative Stage, RS: Reproductive Stages, and the numbers alongside the trait abbreviations indicate the week of data collection.

**Table 1.**
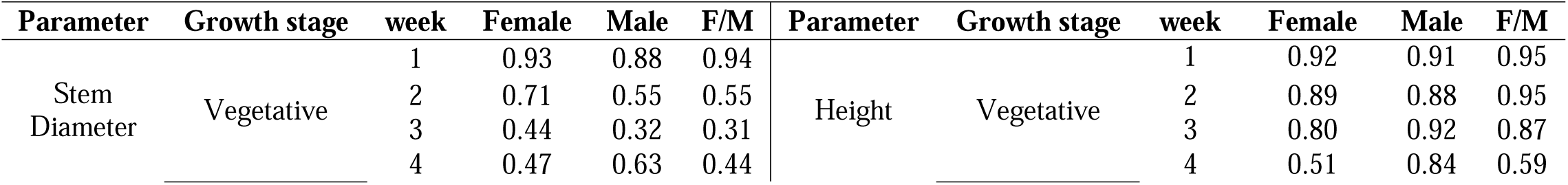
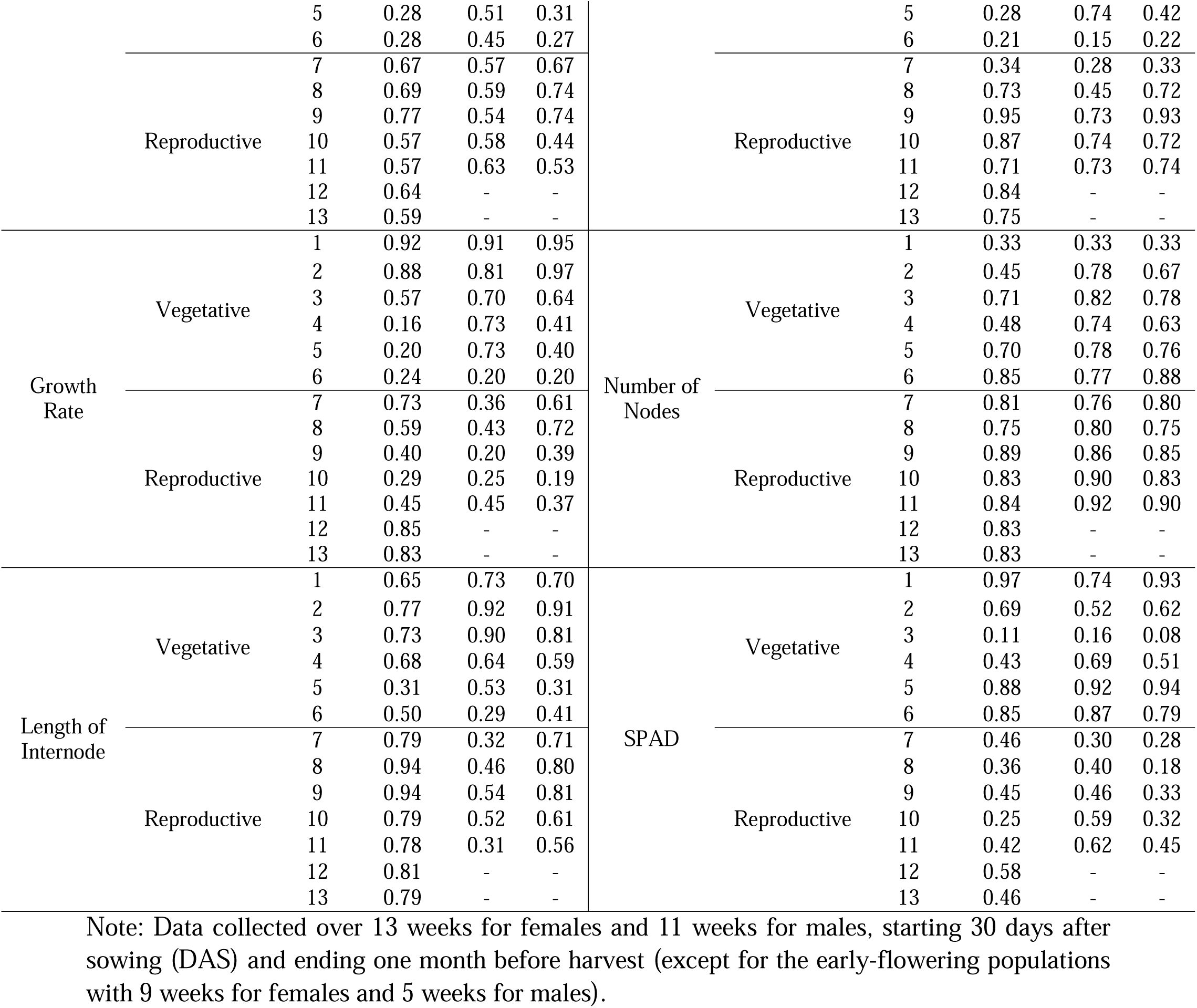
Broad-sense heritability (H^2^) for six key morphophysiological traits in 25 native cannabis populations, assessed weekly during both vegetative (VS) and reproductive stages (RS) in females, males, and both together.

#### i. The analytical temporal framework

The trend of variations in SD, H, GR, NN, LI-MT, and SPAD in female and male plants across different populations (Figs. 3 and 4) exhibited a distinct growth pattern, with notable variations in vigor observed during weekly measurements throughout the cultivation period in both female and male plants. This variance was minimal in the initial weeks of data collection but became more pronounced each week. We found that the highest and lowest SD throughout the growth period were not specific to a particular population (Fig. 3A and B) and varied among different populations over different weeks, except for weeks 8, 9, 10, and 11 in male plants where the population IR4232 consistently had the highest SD, and similarly in female plants where the population IR3776 had the highest SD in weeks 10, 11, 12, and 13. In addition, the range in maximum and minimum SD across different weeks demonstrated a diverse spectrum of population variations. For instance, in the first week of measurements, the SD in male and female plants was 0.3 mm, while in the final week of measurements, the range extended to 3.9 mm in female plants and 2.3 mm in male plants.

The H measurements across different populations (Fig. 3C and D) revealed that the IR5777 population consistently exhibited the highest H measurements from weeks 6 to 13 in both female and male plants. Moreover, the male plants of the IR6522 population also showed the highest H measurements in weeks 9, 10, and 11, like the IR5777 population, indicating a growth surge for this population after week 7. The investigation of weekly GR changes indicated that all populations had a growth peak (maximum GR) followed by a decrease in GR (Fig. 3E and F). This peak occurred in different weeks for different populations, and based on considering this growth peak, we clustered populations in different groups. Accordingly, female plants of different populations were classified into 4 clusters, meaning that populations showed the growth peak in 4 different weeks, including weeks 6, 7, 8, and 9 of data collection (Fig. 3E). Conversely, male plants of different populations showed maximum GR in week 5, 7, and 9 (Fig. 3F).

In the first week of data collection, all populations exhibited the same NN with three nodes (Fig. 4A and B). However, by the fifth week, diversity among populations had increased, with female plants showing a range of three nodes and male plants with a range of four nodes. Aside from minor fluctuations during different weeks, populations consistently followed an increasing trend. The patterns of LI-MT revealed a diverse range of values (Fig. 4C and D). In different populations, despite minor fluctuations during various data collection weeks and the use of different algorithms, the LI-MT range has increased in some weeks; for example, in female plants, weeks 7, 8, and 11 (Fig. 4C), and in male plants, weeks 4, 8, and 9 (Fig. 4D). Additionally, populations IR2385 and IR4457 had the highest LI with a noticeable difference compared to other populations, starting from weeks 7 and 8, where they were closer to other populations before that.

Finally, exploration of SPAD values and patterns across all population (Fig. 4E and Fig. 4F) showed that different populations exhibited a simultaneous and similar doubling of their SPAD values after the second week of data collection. Furthermore, except for minor fluctuations in some weeks in certain populations, the SPAD range indicated a consistent trend from the first to the last week of data collection (ranging between 6-15 in females and 6-18 in males). Additionally, populations IR2385 and IR4457 consistently had the highest SPAD values across different weeks.

#### ii. Changes between vegetative (VS) and reproductive stages (RS)

The frequency distribution of the weekly trends of six parameters (SD, H, GR, NN, LI-MT, and SPAD) in female and male plants across different populations showed that the traits in the VS are mostly skewed to the left (Fig. 5E and F for GR and Fig. 5K and L for SPAD), while in the RS, it predominantly indicated a normal distribution. It is important to note that in the RS, some traits (i.e., SD, H, and NN) exhibited a wider range of variation than the VS.

In the VS, the ranges for both parameters SD (Fig. 5A and B) and H (Fig. 5C and D) in female and male plants were similar, ranging from 0.45 to 6 mm and 4 to 44 cm, respectively. However, the SD ranges also widened during the RS, with female plants ranging from 5 to 16 mm and male plants ranging from 6 to 13 mm (Fig. 5A and B). Meanwhile, during the RS, the ranges for H expanded significantly, showing a range from 40 to 190 cm in female plants and 45 to 160 cm in male plants (Fig. 5C and D).

Over the transition from VS to RS, some traits showed overlap between both stages, such as GR (Fig. 5E and F) and SPAD (Fig. 5K and 5L). Specifically, the range of GR in females were from 0.125 to 2.8 cm/day and 0.9 to 4.5 cm/day during VS and RS, respectively, while in males it ranged from 0.08 to 3.2 cm/day during VS and 1.15 to 5.2 cm/day during RS. In addition, the SPAD index is depicted differently due to a notable gap in the histogram during VS, which has been split into two distinct frequency distributions. Interestingly, both the value and range of the SPAD index were similar in males and females in both VS and RS.

#### iii. Growth pattern differences between female and male plants

Although various patterns demonstrated irregular dynamics in six examined traits across different populations, they still possessed a general and robust framework capable of expressing the growth pattern for both genders. To achieve this, we visualized the trends of the examined traits over 13 weeks for female plants and 11 weeks for male plants by averaging data from all different populations (Fig. 6). Additionally, we defined equations for predicting the values of each trait and placed them in the corresponding figure for each trait, enabling estimation of each trait’s value in the specified week (*x*) by substituting the week number into the relationship *z* = (*x* - 7)/3.89. Parameters SD, H, and NN exhibited similar linear relationships with values of 0.98, 0.95, and 0.99 (Fig. 6A, B and D). On the other hand, parameters GR, LI-MT, and SPAD followed polynomial patterns with a degree of 6 for GR and cubic (i.e. degree of a variable as 3) for the other two, with R^2^ values of 0.99, 0.97, and 0.92 (Fig. 6C, E and F).

As indicated in Fig. 6, the weeks corresponding to the VS and RS are determined based on the photoperiod switch (switch-off), when the length of the day begins to shorten. Male and female plants exhibited similar growth patterns in various traits until the photoperiod switch. However, the differences were observed after the critical photoperiod between male and female plants. The SD showed a growth spurt after the seventh week (SD_RS7), followed by a gradual increase until week 11 (SD_RS11). Subsequently, in female plants, there is a renewed upward trend, with an average increase from 11.5 mm in SD_RS11 to 14 mm in week 13 (SD_RS13) (Fig. 6A). The H increase has continued with a steeper slope from week 6 (H_VS6) onward (Fig. 6B), while the NN has increased steadily with a constant slope (Fig. 6D).

Up until the third week, the GR remained constant, approximately between 0.2 to 0.3 cm/day (Fig. 6C). However, after that, the GR increased, especially in weeks 7 to 9, reaching its peak. By week 9 (GR_RS9), female plants reached a rate of 3 cm/day, while male plants reached 3.4 cm/day. An interesting point is that after three weeks from the photoperiod switch, the GR began to decrease. From weeks 4 to 8, the LI-MT showed an increasing trend, which continued with minor changes until week 11 (LI_RS11). However, in female plants, this distance decreased from 12 to 7 cm between weeks 11 and 13 (Fig. 6E). The analysis of the SPAD index indicated a 100% increase in chlorophyll content from week 2 to 3. However, after that, it showed minor fluctuations and maintained a constant value until the end of the growth period (Fig. 6F).

#### iv. Broad-sense heritability (H^2^)

The trend of variations of broad-sense heritability (H^2^) was estimated for six morphophysiological traits during different weeks in the VS and RS, for both female and male plants, as well as without considering gender (Table 1). We observed interesting variations in H^2^ over the weeks in different growth stages (VS and RS), along with an analysis of the impact of gender on heritability. Among the four out of six traits examined, including SD, H, GR, and SPAD, the heritability values in the first VS weeks were highest in female and male plants. The SD in the first VS week showed values of 0.93, 0.88, and 0.94 in female and male plants, and both, respectively. Similarly, these values in H and GR in the first VS week were completely similar, with values of 0.92, 0.91, and 0.95 in female and male plants, and both, respectively. Additionally, in week 3, the SPAD index exhibited the lowest heritability, with values of 0.11, 0.16, and 0.08 in female plants, male plants, and both together, respectively. This contrasted with the range of 0.74 to 0.97 observed in the first week in the SPAD index. The trend of H^2^ was different in the NN, with the lowest heritability of 0.33 in the first week, it increased in subsequent weeks, showing a slight fluctuation, and remained almost constant across the trial (ranges between 0.45 to 0.92). Additionally, this value showed the highest heritability in the LI-MT, with 0.94 in female plants in weeks 8 and 9, and 0.92 in male plants in week 2.

The heritability values of the examined traits demonstrated that they were influenced by the photoperiod switch during the weeks before and after the switch (transition from VS to RS between weeks 5-8). In SD, this value decreased in weeks 5 and 6 (0.28 in female plants and 0.45 in male plants) and increased in weeks 7 and 8 (after the switch). The H and GR exhibited a significant decrease in week 6 (before the switch), while in the SPAD index, this value decreased by 46% in female plants and 65% in male plants in week 7 (after the switch) compared to week 6.

Additionally, the correlation analysis of six morphophysiological traits over 13 consecutive weeks in female plants (Fig. S1) and 11 weeks in male plants (Fig. S2) revealed that, in male plants, there was a very strong positive correlation among these six traits during weeks 6 to 11. In contrast, during the same period (weeks 6–11), these traits exhibited strong negative correlations with GR, H, NN, and LI-MT in weeks 3, 4, and 5. In female plants, the NN trait showed strong negative correlations with five other traits across various weeks (weeks 2–9), while positive correlations among these traits were more pronounced from weeks 8 to 13.

### 3.2 A framework to identify key biological features of early, medium, and late flowering plants in cannabis

In this study, we propose a framework that integrates three machine learning FS techniques to identify key features highly correlated with flowering time in cannabis. Within this framework, two techniques, MI and RFE, were independently utilized alongside the SVM classification model (i.e., MI-SVM and RFE-SVM). Additionally, the RF model was incorporated as an embedded FS technique to enhance understanding of the correlations and classifications of flowering time (i.e., early, medium, and late) in various phenological stages in cannabis.

The MI model ranked 128,725 out of 234,002 features based on their potential for achieving the highest accuracy in categorizing a panel of 145 cannabis landraces, including male and female plants, into three flowering time categories: early, medium, and late. The SVM model was used as a wrapping to utilize the selected MI-ranked feature list. Initially, the SVM achieved a weighted accuracy of 86.2% using the top four MI-ranked features. As illustrated in Fig. S3A, the accuracy performance of the SVM classifier showed a gradual decline as more features were added. Notably, while three features achieved 86.2% accuracy, we opted for the four features due to their higher frequency of occurrence as key predictors (Table S3). For the RFE-SVM approach, we started with 1,000 initial features, culminating in a remarkable 96.6% weighted accuracy utilizing only 53 features through the RFE-SVM approach (Fig. S3B and Table S4). Furthermore, we optimized the RF model by exploring various numbers of trees, ranging up to 1,000, to determine the optimal configuration. Remarkably, with 197 trees, the model achieved a performance level of 65.5% (Fig. S3C and Table S5).

The evaluation of different techniques included a comparison based on precision, recall (sensitivity), specificity, balanced accuracy, F1-score and ROC AUC score, as summarized in Table 2. The heatmap of the confusion matrix was generated to compare different FS techniques in predicting the number of early, medium, and late flowering accessions (Fig. S4). Furthermore, ROC AUC curves for FPR and TPR are illustrated in Fig. S5.

**Table 2.** Comparison of evaluation metrics among different feature selection (FS) techniques: MI-SVM, RFE-SVM, and RF models. The metrics include precision, recall (sensitivity), specificity, Balanced Accuracy, F1-score, and ROC AUC.

| Technique | MI-SVM | RFE-SVM | RF |
| --- | --- | --- | --- |
| Number of Features | 4 | 53 | 2840 (197 Trees) |
| Precision | 0.862 | 0.968 | 0.666 |
| Recall (Sensitivity) | 0.862 | 0.966 | 0.655 |
| Specificity | 0.879 | 0.971 | 0.699 |
| Balanced Accuracy | 0.87 | 0.969 | 0.677 |
| F1-Score | 0.862 | 0.965 | 0.654 |
| ROC AUC | 0.917 | 0.99 | 0.802 |

In total, 4 features were common across all methods (Fig. 7). Additionally, 24 features were identified by at least two FS techniques. Of the 53 features identified by RFE-SVM, 4 overlapped with MI-SVM and 28 with RF. On the other hand, MI-SVM and RF shared 4 common features (Table 3).

**Fig. 7.**
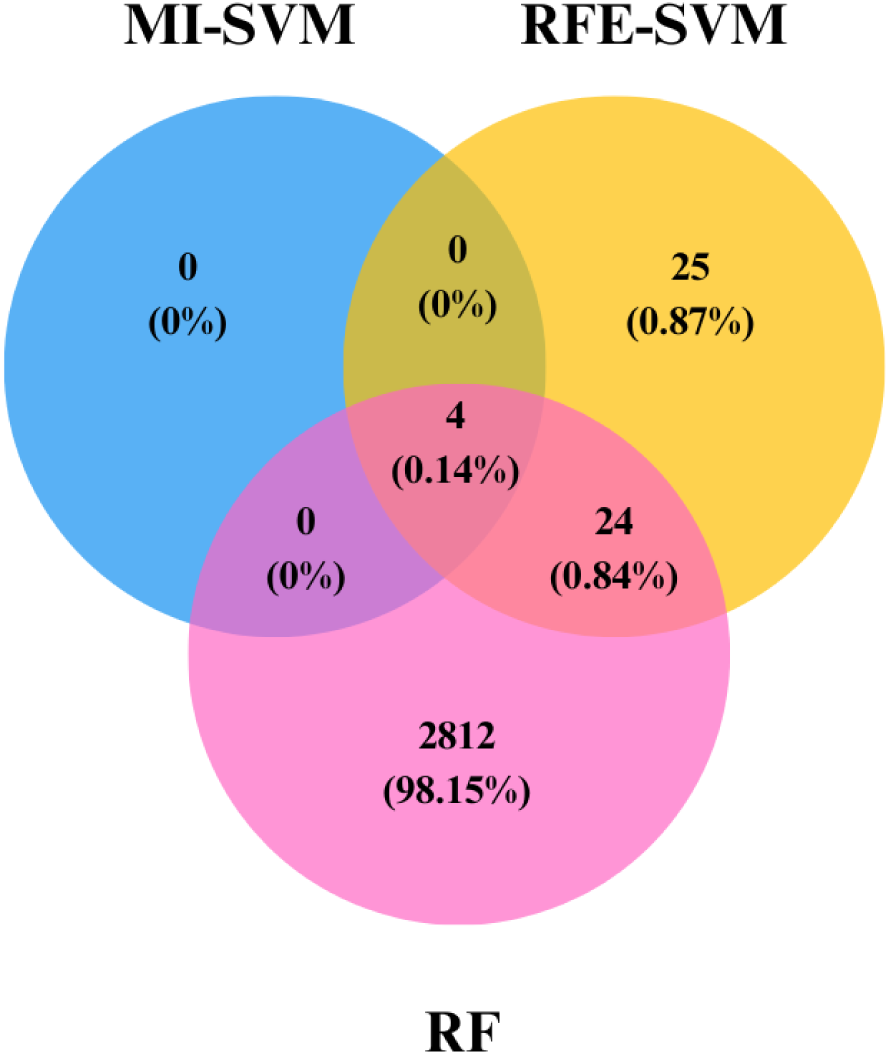
The number of features in each model and the common features across all approaches that differentiate cannabis flowering classes.

**Table 3.** List of features common between all and/or at least two selection techniques.

| Features common between all selection techniques | Features common between at least two selection techniques |  |  |  |  |
| --- | --- | --- | --- | --- | --- |
| LI_MT_RS8 | H_RS11 | SPAD_RS11 | LI_MT_VS6 | GR_RS8 | SD_RS11 |
| LI_MT_RS11 | LI_MT_RS7 | NN_RS10 | ChrX_44207464 | GR_RS7 | GR_RS10 |
| LI_MT_RS9 | SPAD_RS10 | H_RS10 | ChrX_44209294 | H_RS7 | Chr01_19732617 |
| LI_MT_RS10 | H_RS8 | GR_RS9 | Chr06_10601607 | SD_RS7 | SPAD_VS6 |
|  | H_RS9 | SD_RS9 | NN_RS11 | SD_RS8 |  |

In the final model, an SVM classification model was constructed using three feature sets based on their highest accuracy, performance and minimal number of features. The three feature sets included: a) the four features common to all methods to evaluate our candidate markers reported by all techniques (Table 3); b) the 28 features identified by at least two methods, including the 4 features common to all (Table 3); and c) the 53 features selected by RFE-SVM, which provided insights with 31 morphophysiological features and 22 unique SNP sites on different chromosomes (Table S4).

The model achieved an accuracy score of 86.2% and ROC AUC score of 0.917 with four features of features set (a), an accuracy score of 86.2% and ROC AUC score of 0.955 with 28 features of features set (b) and with the 53 features of features set (c) achieved the highest accuracy and ROC AUC scores of 96.6% and 0.994, respectively. Other evaluation measures have also been computed (Table 4), along with the heatmap of the confusion matrix and ROC AUC curves shown for feature sets a and c in Fig. 8 and for feature set b in Fig. S6.

**Fig. 8.**
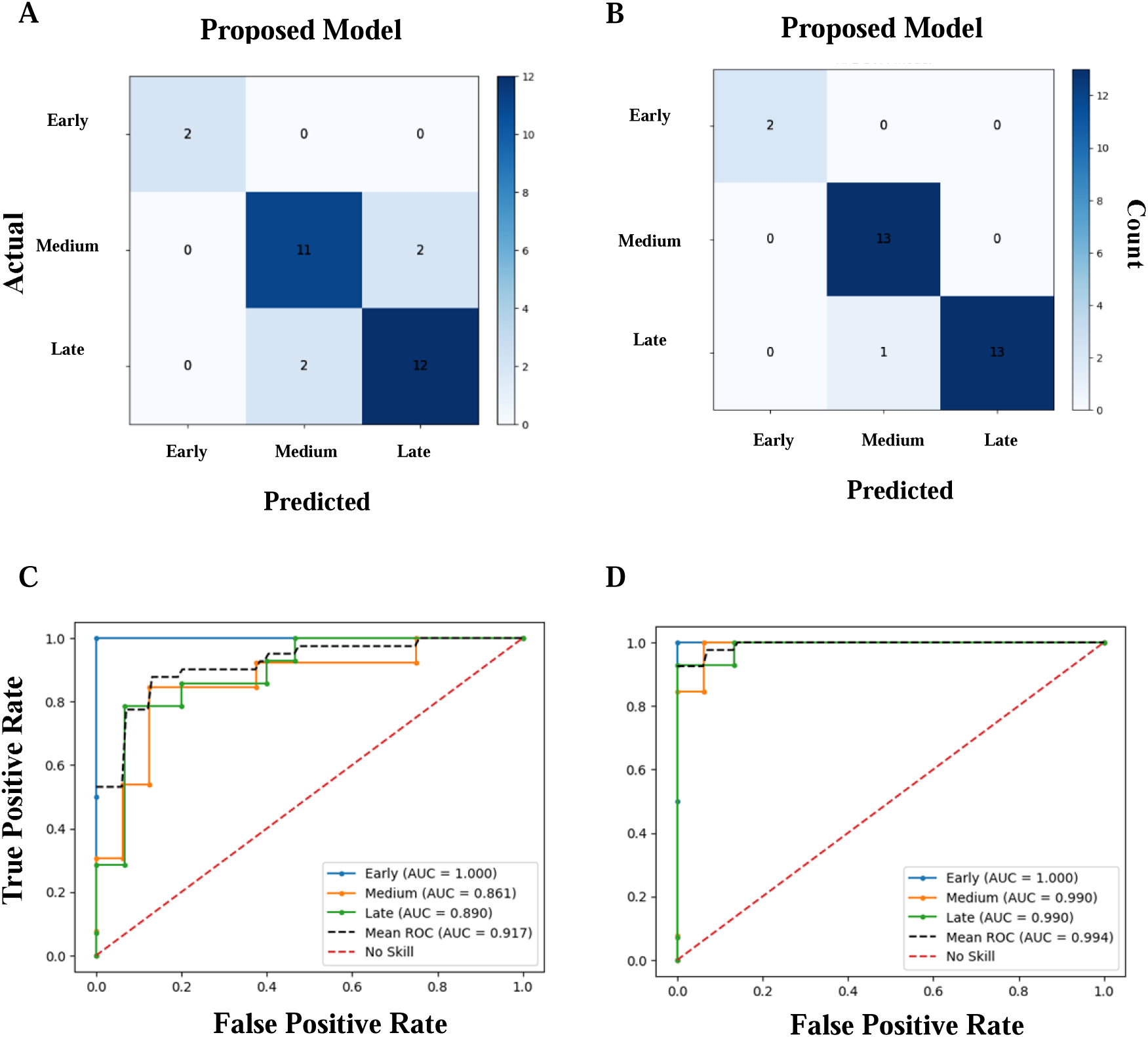
Confusion matrix heatmap using the proposed model of (**A**) features common between all selection techniques (feature = 4), and (**B**) features selected with RFE-SVM (feature = 53). ROC AUC validation used the proposed model of (**C**) features common between all selection techniques, and (**D**) features selected with RFE-SVM that differentiate cannabis flowering classes. ROC AUC and confusion matrix are based on the test set, representing 20% of the total data.

**Table 4.**
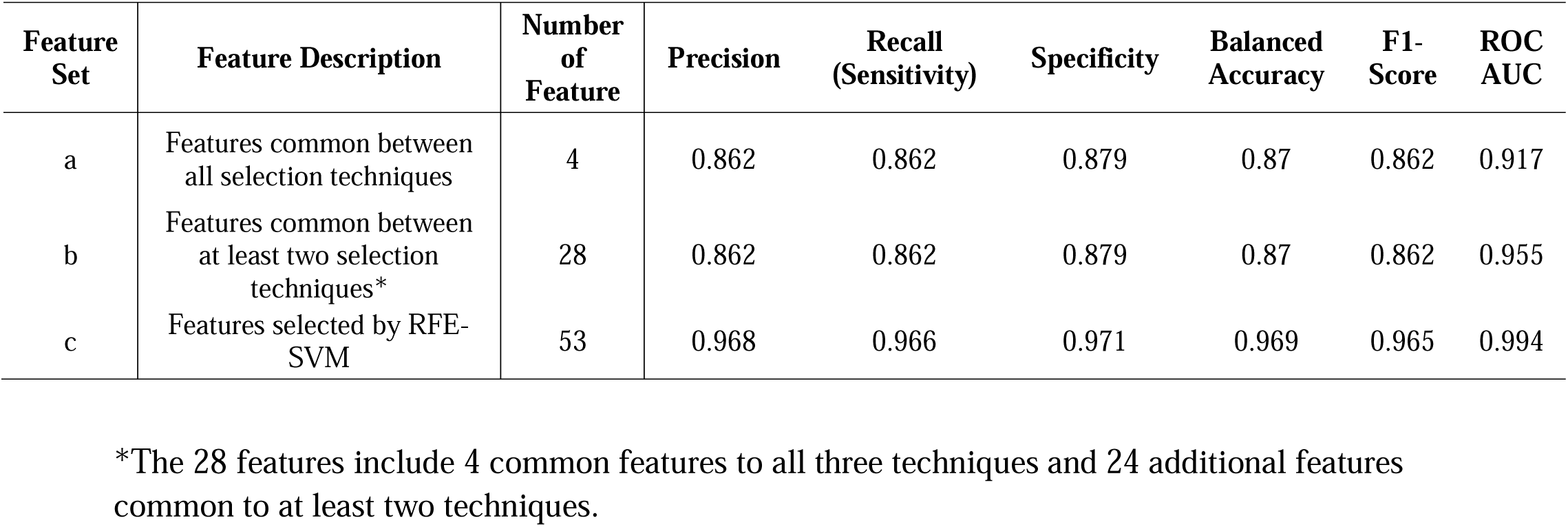
Evaluation metrics for the proposed model based on features common between all, at least two selection techniques and features selected with RFE-SVM that differentiate cannabis flowering classes.

| Feature Set | Feature Description | Number of Feature | Precision | Recall (Sensitivity) | Specificity | Balanced Accuracy | F1-Score | ROC AUC |
| --- | --- | --- | --- | --- | --- | --- | --- | --- |
| a | Features common between all selection techniques | 4 | 0.862 | 0.862 | 0.879 | 0.87 | 0.862 | 0.917 |
| b | Features common between at least two selection techniques* | 28 | 0.862 | 0.862 | 0.879 | 0.87 | 0.862 | 0.955 |
| c | Features selected by RFE-SVM | 53 | 0.968 | 0.966 | 0.971 | 0.969 | 0.965 | 0.994 |

\*The 28 features include 4 common features to all three techniques and 24 additional features common to at least two techniques.

To visualize the distribution of features across early, medium, and late flowering classes, boxplots were generated for each of the three feature sets (a, b, and c) identified by the ML framework (Figs. S7–S9). These boxplots highlight the variability and separation of feature values such as internode length and genetic markers (SNPs) on chromosomes 01, 08, 09, and X. The early-flowering group consistently showed distinct separation in feature distributions across all feature sets, reflecting their unique developmental trajectory. In contrast, medium- and late-flowering groups exhibited overlapping distributions for traits like SD and H during the reproductive stage, indicating closer phenotypic and genetic similarities.

To evaluate the diagnostic potential of the introduced markers in the three feature sets (a, b, and c) within the proposed model, ROC AUC analysis was conducted using 5-fold cross-validation to distinguish between the three flowering classes in 145 native cannabis accessions (Fig. 9). The results are presented in two ways: the AUC value of each marker in identifying each flowering class (Figs. S10, S12, and S14 for feature sets a, b, and c, respectively) and the average AUC value of each marker in identifying all three flowering classes (Figs. S11, S13, and S15). The results indicated that most markers exhibit strong capability in identifying early-flowering plants. However, there were fewer markers with an accuracy above 70% in distinguishing between medium- and late-flowering plants, suggesting that the model achieves high accuracy by correlating these features in the proposed model, with average AUC values of 0.917, 0.955, and 0.994 for feature sets a, b, and c, respectively.

**Fig. 9.**
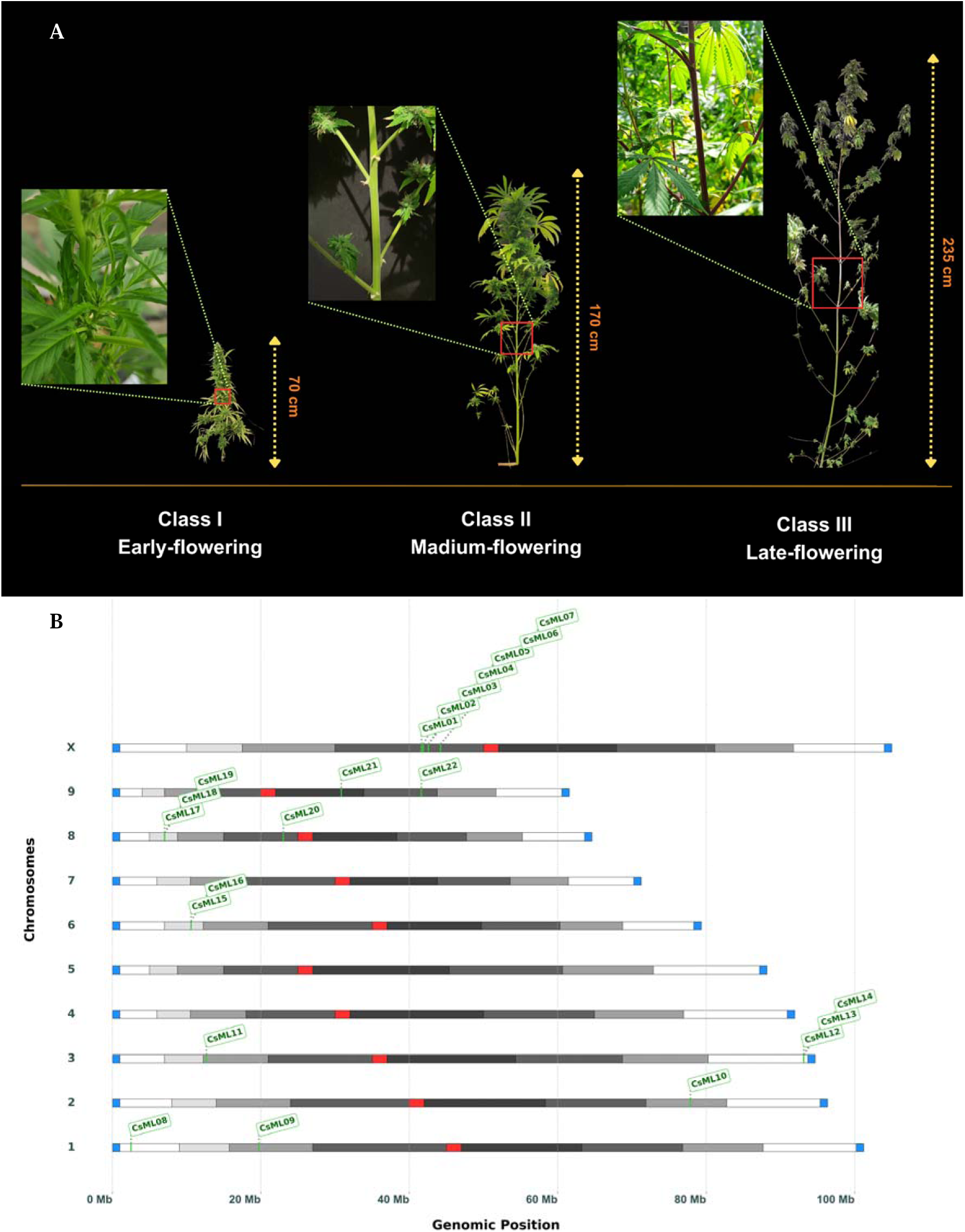
Key morpho-physiological and genetic features differentiate cannabis flowering classes. (**A**) Morphological variations in internode length (LI_MT_RS) were identified by all methods as potential markers. (**B**) Distribution of 22 candidate SNP markers on different chromosomes, highlighting their role in flowering time differences among the three cannabis flowering classes.

Feature set c, which contains 53 candidate markers identified by the RFE-SVM, had the highest AUC value of 0.994 among the three selected feature sets. This set included 31 morphophysiological markers recorded from weeks 4 to 11, covering the six morphophysiological traits examined in this study. Specifically, in weeks 4 and 5, only the NN trait was included; in week 6, two traits, LI_MT and SPAD, were added; in weeks 7, 8, and 9, all traits except NN were included; and in weeks 10 and 11, all six traits (NN, LI_MT, SPAD, SD, H, and GR) were considered. Additionally, this feature set introduced 22 candidate genetic markers (SNPs) distributed across the genome, including the X sex chromosome with 7 SNPs, chromosome 01 with 2 SNPs, chromosome 02 with 1 SNP, chromosome 03 with 4 SNPs, chromosome 06 with 2 SNPs, chromosome 08 with 4 SNPs, and chromosome 09 with 2 SNPs (Figs. 9, S14, S15 and Table S4). These SNPs were located within 11 haplotype blocks identified via LD analysis, with block sizes ranging from 0.008 to 11.4 kb and representative pairwise LD (*r²*) values between 0.75 and 0.96 (Fig. S16 and Table S6). Some of these markers were positioned within or adjacent to annotated genes (Table S6).

## 4. Discussion

This study introduces an integrative framework to classify flowering time in *Cannabis sativa* using longitudinal morphophysiological traits and genomic data. By tracking temporal growth patterns across 145 accessions, we identified early-, medium-, and late-flowering types with high accuracy (96.6%) using a reduced set of 53 features. Unlike Genome-Wide Association Study (GWAS) approaches that rely on single-time-point traits, our method captures dynamic physiological transitions over the full growth cycle. This allows the detection of biologically meaningful patterns that reflect the developmental divergence among flowering groups in cannabis.

The three flowering classes exhibited distinct developmental trajectories, likely shaped by their responses to photoperiod and internal physiological cues. The clear separation of the early-flowering group in feature distributions reflects a pronounced developmental divergence, likely driven by genetic adaptations such as photoperiod insensitivity (Toth et al. 2022; Dowling et al. 2024). This distinct clustering aligns with the day-neutral characteristics of early-flowering landraces, which are adapted to initiate reproduction earlier, potentially optimizing yield in short-season environments (Amaducci et al. 2008; Stack et al. 2021). In contrast, the overlapping feature distributions between medium- and late-flowering groups suggest shared genetic ancestry or less pronounced differentiation in photoperiod-sensitive pathways, complicating their classification (McKernan et al. 2020; Petit et al. 2020a). The dioecious nature of cannabis, coupled with high heterozygosity and the inclusion of both sexes in our diverse landrace panel, further increased the complexity of the classification space. Despite these challenges, the ML framework effectively identified markers combinations that achieved high classification accuracy. This demonstrates the framework’s ability to resolve complex trait interactions in genetically diverse populations, providing robust insights for breeding programs targeting tailored flowering times (Montesinos-López et al. 2018; Cherif et al. 2023; Mora-Poblete et al. 2023; Chavhan et al. 2024; Farooq et al. 2024).

Beyond classifying flowering types, our study provides insights into the weekly dynamics of morphophysiological traits and their genetic control. We observed varying broad-sense heritability (H^2^) for traits across different weeks, with notable shifts influenced by environmental cues, particularly the photoperiod switch-off (weeks five to eight) during the vegetative-to-reproductive phase transition (Grassi and McPartland 2017; Naim-Feil et al. 2021, 2023; Babaei et al. 2022). For instance, a significant decrease in H^2^ for the SPAD index in the third week coincided with a 100% increase in SPAD across all genotypes (Chaimala et al. 2023; Chen et al. 2023; Tang et al. 2024). This synchronized increase in chlorophyll production across all genotypes in early growth stages enhances photosynthesis and metabolism (Morales et al. 2020; Cackett et al. 2022; Shi et al. 2022b). This uniform change suggests an environmental factor acted as a cue, leading to similar physiological changes in cannabis plants of different genotypes during the juvenile growth stage (Amaducci et al. 2008; Babaei et al. 2024). Similarly, while plant height and stem diameter showed diverse growth patterns reflecting varied vigor, they also exhibited different responses to short-day light induction. Notably, stem diameter ceased growth following the switch-off, except for female plants, which showed an increased trend in diameter from five weeks after the switch-off until the end of the growth period, similar to previous reports (Obeso, 2002; Pantin et al., 2012; Naim-Feil et al., 2021).

The stability of the SPAD index from the third week of the vegetative stage until the end of the growth period, as demonstrated in our study, suggests plants maintain photosynthesis relatively unaffected by the switch-off, even as other growth indices are influenced. This indicates the plant perceives day length change as a physiological cue, regulating the transition to the reproductive phase by adjusting photosynthetic product partitioning, a concept consistent with photoperiodism’s role in plant development (Adams and Langton 2005; Song et al. 2015; Gawarecka and Ahn 2021; Chaimala et al. 2023; Gendron and Staiger 2023).

Our analysis of temporal morphophysiological dynamics revealed notable changes in the distribution and range of traits between vegetative and reproductive stages. In early growth, plants from different populations exhibited similar trait values; however, variability increased over time, consistent with findings by. This expanding divergence was critical for differentiating flowering-time classes using ML models. Although the observed range of growth rates between growth phases differed from those reported by Naim-Feil et al. (2021) —likely due to our use of genetically diverse seedlings as opposed to clonal cuttings (Sultan 2000; Li et al. 2022; Godwin et al. 2024)—the general trend of dynamic trait behavior remained evident.

The observed divergence in morphophysiological changes, along with transitions across phenological stages, are orchestrated by a broad network of genes interacting with environmental factors such as day length and temperature (Andrés and Coupland 2012; Hill and Li 2016; Babaei et al. 2022). Cannabis plants synchronize biological activities with environmental rhythms by regulating their internal circadian phase, ensuring optimal performance across diurnal and seasonal cycles (Webb 2003; Tan and Swain 2006; Hüve et al. 2007; Harmer 2010; Gottlieb 2019; Webb et al. 2019; Steed et al. 2021). This alignment affects diverse functions, including flower emergence, and enhances plant fitness (Michael et al. 2003; Dodd et al. 2005).

The dynamic interactions among morphophysiological traits, as revealed by our correlation analyses, strongly support this. Traits such as growth rate, height, and internode length showed interconnected responses across specific weeks, reflecting how developmental and environmental rhythms, coordinated by circadian and hormonal pathways, modulate these characteristics and collectively shape the trajectory towards flowering. Understanding these interdependencies at critical weekly intervals is key to dissecting the complex physiology leading to different flowering times (Turck et al. 2008; Petit et al. 2020a; Alter et al. 2024).

The SNPs identified in this study revealed strong genetic signals associated with early flowering, offering new insights into the complex genetic control of photoperiod insensitivity in cannabis. Several markers identified in this study showed exceptionally high classification performance (ROC AUC 96–100%) and were uniquely present in early-flowering accessions, suggesting their potential functional relevance. Among them, three closely linked SNPs on chromosome 8—*CsML17*, *CsML18*, and *CsML19*—were located within a shared ∼3 kb haploblock exclusive to early-flowering landraces. Given its unique structure and strong discriminatory power, we propose this region as a novel early-flowering locus, designated *Autoflower3*(*CsFT3*).

In cannabis, the photoperiod pathway is a major determinant of flowering, with genes such as *CsFT2*, a member of the PEBP gene family (*FLOWERING LOCUS T* homolog), linked to photoperiod sensitivity and flowering induction (Turck et al. 2008; Steel et al. 2023; Dowling et al. 2024). Previous studies have identified early-flowering loci in cannabis, such as *Autoflower1* (*Early1*) on chromosome 1 in high-cannabinoid accessions (Toth et al. 2022) and *Autoflower2* (*FT1*) at the distal end of chromosome 8 in hemp cultivars (Dowling et al. 2024). In contrast, the markers comprising *AutoFlower3* (*CsFT3*) (*CsML17*, *CsML18*, and *CsML19*), as identified in our study, are located in a different genomic region. This divergence may reflect the influence of structural variation, gene duplication, or population-specific haplotypes that lead to the identification of distinct but functionally related loci across studies (Spielmann et al. 2018; Chen et al. 2024; Michael et al. 2024). Additional high-confidence markers were also identified on chromosomes X, 1, and 9, including *CsML03–07* (two different haploblocks on chromosome X, *LOC115711146*), *CsML09* (chr1), and *CsML21* (chr9, *LOC115722557*, *CircadianFloweringLocus1* (Cs*CFL1*)), all contributing to early-flowering classification. These findings support the hypothesis that early flowering in cannabis arises from a polygenic and structurally diverse genetic architecture. This complexity is further reinforced by pangenomic studies showing that structural variations (SVs)—such as duplications, inversions, and translocations—and transposable elements (TEs) can modulate gene regulation and contribute to functional diversity across accessions, as also seen in *Humulus lupulus* (De Oliveira et al. 2020; Loiseau et al. 2020; Dowling et al. 2021, 2024; Ferraj et al. 2023; Michael et al. 2024).

Hormonal pathways also contribute significantly to flowering regulation, with gibberellins (GA3) and auxins (IAA) playing central roles in the vegetative-to-reproductive transition (Chandler 2011; Ajdanian et al. 2023, 2024; Chaudhry et al. 2024). In cannabis, the interplay between hormonal regulation and morphological traits such as internode length, a marker identified in our study, reflects the integration of genetic and hormonal signals (Salentijn et al. 2019; Stack et al. 2021; Ajdanian et al. 2024). Under short-day (SD) conditions, reduced GA3 levels in the shoot apex have been shown to correlate with compact inflorescences and shorter internodes, a characteristic also observed in our study (Alter et al. 2024). Furthermore, exogenous application of GA3 has been reported to mimic long-day (LD) conditions, resulting in elongated internodes and disassembly of condensed inflorescences (Luo et al. 2021; Ajdanian et al. 2024). Similarly, changes in auxin levels under SD conditions may contribute to altered branching patterns and reduced internode elongation, though their precise role requires further investigation (Wei et al. 2022; Alter et al. 2024). Selective breeding has shaped the hormonal regulation and morphology of cannabis, with traits like internode length directly influencing flowering patterns. Fiber cultivars, driven by higher GA3 activity, develop elongated internodes and exhibit delayed flowering, while seed-propagated cultivars, with reduced GA3 levels, show compact growth and earlier flowering transitions (Lalge et al. 2016; Mendel et al. 2020; Wei et al. 2022; Alter et al. 2024).

Developmental timing mechanisms, such as those involving microRNAs like *miR156* and *miR172*, mediate the vegetative-to-reproductive transition by targeting transcription factors in GA3 signaling pathways (Petit et al. 2020a; Steel et al. 2023). These microRNAs have been shown to regulate traits such as internode elongation and flowering in response to environmental and hormonal signals, providing a fine-tuned balance between growth and reproduction (Wu and Poethig 2006; Steel et al. 2023). The interplay of our identified markers with these developmental pathways could help explain the observed diversity in flowering patterns. In summary, our findings highlight the multifaceted genetic control of flowering time in *Cannabis sativa*, with the identified markers potentially influencing photoperiod sensitivity, circadian regulation, hormonal signaling, and epigenetic modifications. These markers may serve as valuable targets for breeding programs aimed at optimizing flowering traits under diverse environmental conditions.

## 5. Conclusion

Understanding flowering time in cannabis is crucial for optimizing cultivation and accelerating breeding programs. Our study employed a novel, data-driven approach integrating longitudinal morphophysiological data, genomic insights, and machine learning (ML) to investigate flowering time diversity in indigenous populations.

Analyzing a comprehensive dataset of over 234k features, we successfully identified 53 key features—including genetic variants and morphophysiological traits—that effectively distinguish between early, medium, and late flowering classes with an impressive 96.6% accuracy. Furthermore, we identified four common morphophysiological features across multiple ML algorithms, achieving 86.2% accuracy in classification.

These findings significantly enhance our understanding of the genetic mechanisms regulating flowering time and reveal critical growth phases. The identified key features and markers support the concept of chronoculture and the development of cannabis as a “smart crop” through AI-driven optimization. Importantly, these markers enable tailored management strategies and early screening in breeding programs, allowing for more efficient selection and acceleration of cultivar development. By integrating these data-driven insights, potentially combined with future advancements like genetic editing, cannabis cultivation can move towards precision strategies for yield optimization and enhanced resilience in diverse environments.

## Abbreviations

ML: Machine Learning
DP: Deep Learning
FS: Feature Selection
SA: Spatial Analysis
DAS: Days After Sowing
SD-VS: Stem Diameter at Vegetative Stage
H-VS: Height at Vegetative Stage
GR-VS: Growth Rate at Vegetative Stage
NN-VS: Number of Nodes on the Main Stem at Vegetative Stage
LI-MT-VS: Length of Internode in the Middle Third of the Main Stem at Vegetative Stage
SPAD-VS: SPAD-based Chlorophyll at Vegetative Stage
SD-RS: Stem Diameter at Reproductive Stages
H-RS: Height at Reproductive Stages
GR-RS: Growth Rate at Reproductive Stages
NN-RS: Number of Nodes on the Main Stem at Reproductive Stages
LI-MT-RS: Length of Internode in the Middle Third of the Main Stem at Reproductive Stages
SPAD-RS: SPAD-based Chlorophyll at Reproductive Stages
VS: Vegetative Stages
RS: Reproductive Stages
MI: Mutual Information
RFE: Recursive Feature Elimination
SVM: Support Vector Machine
RF: Random Forest
SF10I: Start 10% Flowering Time in Individuals (10% of bracts formed)
ROC: Receiver Operating Characteristics
AUC: Areas Under the Curve
FPR: False Positive Rate
TPR: True Positive Rate
CsCFL1: CircadianFloweringLocus1
CsFT3: AutoFlower3
HB: Haplotype Block
LD: Linkage Disequilibrium

## Statements and Declarations

### Author Contributions

Conceptualization: M.B, H.N, H.A, D.T.; Methodology: M.B. (ML/statistical methods, experimental setup, and genomic workflows); Investigation: M.B. (Plant maintenance, seeds and data collection); Data Curation: M.B. (morphophysiological/genomic datasets); Formal Analysis: M.B. (statistics, ML modeling); Software: M.B. (ML implementation); Resources: M.B, H.N, H.A, D.T; Supervision: H.N, D.T.; Funding Acquisition: D.T.; Visualization: M.B.; Writing – Original Draft: M.B.; Writing – Review & Editing: All authors.

### Data Availability

The raw datasets generated during the current study are available. Processed data supporting the findings are publicly accessible via Figshare (DOI: <u>10.6084/m9.figshare.29054192</u>). The custom Python 3.12 codebase, developed using the scikit-learn library for machine learning workflows, is available at https://github.com/Mehdibabaeii/CannaFeatML.

### Ethics approval

Seed collection and cultivation were conducted with ethics approval and license from the anti-narcotics police in Razavi Khorasan. Activities at Université Laval complied fully with Health Canada regulations under license LIC-QX0ZJC7SIP-2021.

## Acknowledgements

The authors wish to thank Ladan Ajdanian for her valuable contribution in data collection and cultivation.

## Reference

Abdelwahab O, Awad N, Elserafy M, Badr E (2022) A feature selection-based framework to identify biomarkers for cancer diagnosis: A focus on lung adenocarcinoma. PLoS One 17:e0269126

Adams SR, Langton FA (2005) Photoperiod and plant growth: a review. J Hortic Sci Biotechnol 80:2–10

Adikaram KKLB, Hussein MA, Effenberger M, Becker T (2015) Data transformation technique to improve the outlier detection power of Grubbs’ test for data expected to follow linear relation. J Appl Math 2015:1–9

Ahmed SA, Ross SA, Slade D, et al (2008) Cannabinoid ester constituents from high-potency Cannabis sativa. J Nat Prod 71:536–542

Ajdanian L, Arouiee H, Jones AMP, et al (2024) Investigating the impact of paclobutrazol and tannic acid on floral development of in vitro-grown cannabis plantlets. Heliyon 10:

Ajdanian L, Babaei M, Arouiee H, et al (2023) Role of ethylene in regulating physiological and molecular aspects of plants under abiotic stress. In: The Role of Growth Regulators and Phytohormones in Overcoming Environmental Stress. Elsevier, pp 113–135

Alejandrino J, Concepcion R, Lauguico S, et al (2020) Visual classification of lettuce growth stage based on morphological attributes using unsupervised machine learning models. In: 2020 IEEE REGION 10 CONFERENCE (TENCON). IEEE, pp 438–443

Alter H, Sade Y, Sood A, et al (2024) Inflorescence development in female cannabis plants is mediated by photoperiod and gibberellin. Hortic Res 11:uhae245

Amaducci S, Colauzzi M, Bellocchi G, Venturi G (2008) Modelling post-emergent hemp phenology (Cannabis sativa L.): theory and evaluation. European Journal of Agronomy 28:90–102

Amaducci S, Pelatti F, Bonatti PM (2005) Fibre development in hemp (Cannabis sativa L.) as affected by agrotechnique: preliminary results of a microscopic study. Journal of industrial hemp 10:31–48

Andrés F, Coupland G (2012) The genetic basis of flowering responses to seasonal cues. Nat Rev Genet 13:627–639

Anwar F, Latif S, Ashraf M (2006) Analytical characterization of hemp (Cannabis sativa) seed oil from different agro ecological zones of Pakistan. J Am Oil Chem Soc 83:323–329

Babaei M, Ajdanian L (2020) Screening of different Iranian ecotypes of cannabis under water deficit stress. Sci Hortic 260:. 10.1016/j.scienta.2019.108904

Babaei M, Ajdanian L, Lajayer BA (2022) Morphological and phytochemical changes of Cannabis sativa L. affected by light spectra. In: New and Future Developments in Microbial Biotechnology and Bioengineering. Elsevier, pp 119–133

Babaei M, Boissinot J, Ajdanian L, Torkamaneh D (2025) Cannabis Genetic Resources. In: The Cannabis Genome. CRC Press, pp 32–41

Babaei M, Nemati H, Arouiee H, Torkamaneh D (2023) Characterization of Indigenous Populations of Cannabis in Iran: A Morphological and Phenological Study. Res Sq

Babaei M, Nemati H, Arouiee H, Torkamaneh D (2024) Characterization of indigenous populations of cannabis in Iran: a morphological and phenological study. BMC Plant Biol 24:151. 10.1186/s12870-024-04841-y

Balant M, Garnatje T, Vitales D, et al (2024) Intra leaf modeling of Cannabis leaflet shape produces leaf models that predict genetic and developmental identities. New Phytologist

Barcaccia G, Palumbo F, Scariolo F, et al (2020) Potentials and challenges of genomics for breeding cannabis cultivars. Front Plant Sci 11:573299

Barrett JC, Fry B, Maller J, Daly MJ (2005) Haploview: analysis and visualization of LD and haplotype maps. Bioinformatics 21:263–265

Belgiu M, Drăguţ L (2016) Random forest in remote sensing: A review of applications and future directions. ISPRS journal of photogrammetry and remote sensing 114:24–31

Braich S, Baillie RC, Spangenberg GC, Cogan NOI (2020) A new and improved genome sequence of Cannabis sativa. GigaByte 2020:

Browning BL, Browning SR (2016) Genotype imputation with millions of reference samples. The American Journal of Human Genetics 98:116–126

Cackett L, Luginbuehl LH, Schreier TB, et al (2022) Chloroplast development in green plant tissues: the interplay between light, hormone, and transcriptional regulation. New Phytologist 233:2000–2016

Cain MK, Zhang Z, Yuan K-H (2017) Univariate and multivariate skewness and kurtosis for measuring nonnormality: Prevalence, influence and estimation. Behav Res Methods 49:1716–1735

Cannoy DC (2015) Green Gold-a Cannabis Sativa L. Lucis Suitability Analysis for West Virginia

Cao S, Luo X, Xu D, et al (2021) Genetic architecture underlying light and temperature mediated flowering in Arabidopsis, rice, and temperate cereals. New Phytologist 230:1731–1745

Carpentier C, Mulligan K, Laniel L, et al (2012) Cannabis production and markets in Europe. Publ. Office of the Europ. Union

Chaimala A, Jogloy S, Vorasoot N, et al (2023) The variation of relative water content, SPAD chlorophyll meter reading, stomatal conductance, leaf area, and specific leaf area of Jerusalem artichoke genotypes under different durations of terminal drought in tropical region. J Agron Crop Sci 209:12–26

Chandler JW (2011) The hormonal regulation of flower development. J Plant Growth Regul 30:242–254

Chaudhry A, Chen Z, Gallavotti A (2024) Hormonal influence on maize inflorescence development and reproduction. Plant Reprod 1–15

Chavhan RL, Hinge VR, Wankhade DJ, et al (2024) Bioinformatics for Molecular Breeding and Enhanced Crop Performance: Applications and Perspectives. Bioinformatics for Plant Research and Crop Breeding 21–74

Chen B, Huang G, Lu X, et al (2023) Prediction of vertical distribution of SPAD values within maize canopy based on unmanned aerial vehicles multispectral imagery. Front Plant Sci 14:1253536

Chen X, Jeong JC (2007) Enhanced recursive feature elimination. In: Sixth international conference on machine learning and applications (ICMLA 2007). IEEE, pp 429–435

Chen Y, Khan MZ, Wang X, et al (2024) Structural variations in livestock genomes and their associations with phenotypic traits: a review. Front Vet Sci 11:1416220

Cherif E, Feilhauer H, Berger K, et al (2023) From spectra to plant functional traits: Transferable multi-trait models from heterogeneous and sparse data. Remote Sens Environ 292:113580

Cingolani P, Platts A, Wang LL, et al (2012) A program for annotating and predicting the effects of single nucleotide polymorphisms, SnpEff: SNPs in the genome of Drosophila melanogaster strain w1118; iso-2; iso-3. Fly (Austin) 6:80–92

Crocq M-A (2020) History of cannabis and the endocannabinoid system. Dialogues Clin Neurosci 22:223–228. 10.31887/DCNS.2020.22.3/mcrocq

Danecek P, Auton A, Abecasis G, et al (2011) The variant call format and VCFtools. Bioinformatics 27:2156–2158

De Meijer EPM, Hammond KM (2005) The inheritance of chemical phenotype in Cannabis sativa L.(II): cannabigerol predominant plants. Euphytica 145:189–198

De Oliveira R, Rimbert H, Balfourier F, et al (2020) Structural variations affecting genes and transposable elements of chromosome 3B in wheats. Front Genet 11:891

de Ronne M, Lapierre É, Torkamaneh D (2024) Genetic insights into agronomic and morphological traits of drug-type cannabis revealed by genome-wide association studies. Sci Rep 14:9162

Dodd AN, Salathia N, Hall A, et al (2005) Plant circadian clocks increase photosynthesis, growth, survival, and competitive advantage. Science (1979) 309:630–633

Dowling CA, Melzer R, Schilling S (2021) Timing is everything: the genetics of flowering time in Cannabis sativa. Biochem (Lond) 43:34–38

Dowling CA, Shi J, Toth JA, et al (2024) A FLOWERING LOCUS T ortholog is associated with photoperiod insensitive flowering in hemp (Cannabis sativa L.). The Plant Journal 119:383–403

Farooq MA, Gao S, Hassan MA, et al (2024) Artificial intelligence in plant breeding. Trends in Genetics

Ferraj A, Audano PA, Balachandran P, et al (2023) Resolution of structural variation in diverse mouse genomes reveals chromatin remodeling due to transposable elements. Cell Genomics 3:

Flores-Sanchez IJ, Verpoorte R (2008) Secondary metabolism in cannabis. Phytochemistry reviews 7:615–639

Gawarecka K, Ahn JH (2021) Isoprenoid-derived metabolites and sugars in the regulation of flowering time: Does day length matter? Front Plant Sci 12:765995

Gendron JM, Staiger D (2023) New horizons in plant photoperiodism. Annu Rev Plant Biol 74:481–509

Godwin A, Pieralli S, Sofkova-Bobcheva S, et al (2024) Comparing vegetative growth patterns of cultivated (Daucus carota L. subsp. sativus) and wild carrots (Daucus carota L. subsp. carota) to eliminate genetic contamination from weed to crop. J Agric Food Res 16:101213. 10.1016/j.jafr.2024.101213

Gottlieb D (2019) Agro-chronobiology: Integrating circadian clocks/time biology into storage management. J Stored Prod Res 82:9–16

Grassa CJ, Wenger JP, Dabney C, et al (2018) A complete Cannabis chromosome assembly and adaptive admixture for elevated cannabidiol (CBD) content. BioRxiv 458083

Grassi G, McPartland JM (2017) Chemical and morphological phenotypes in breeding of Cannabis sativa L. Cannabis sativa L-Botany and Biotechnology 137–160

Harmer S (2010) Plant Biology in the Fourth Dimension. Plant Physiol 154:467–470. 10.1104/pp.110.161448

Hill CB, Li C (2016) Genetic Architecture of Flowering Phenology in Cereals and Opportunities for Crop Improvement. Front Plant Sci 7:

Hüve K, Christ MM, Kleist E, et al (2007) Simultaneous growth and emission measurements demonstrate an interactive control of methanol release by leaf expansion and stomata. J Exp Bot 58:1783–1793

Ibrar D, Ahmad R, Hasnain Z, et al (2023) Epigenetic based control of flowering and seed development in plants: A review. Plant Breeding 142:732–744

Ii APG (2003) An update of the Angiosperm Phylogeny Group classification for the orders and families of flowering plants: APG II. Botanical journal of the Linnean Society 141:399–436

Jakkula V (2006) Tutorial on support vector machine (svm). School of EECS, Washington State University 37:3

Khalili E, Kouchaki S, Ramazi S, Ghanati F (2020) Machine learning techniques for soybean charcoal rot disease prediction. Front Plant Sci 11:590529

Kovalchuk I, Pellino M, Rigault P, et al (2020) The genomics of Cannabis and its close relatives. Annu Rev Plant Biol 71:713–739

Kraskov A, Stögbauer H, Grassberger P (2004) Estimating mutual information. Physical Review E—Statistical, Nonlinear, and Soft Matter Physics 69:066138

Kumari P, Khan S, Wani IA, et al (2022) Unravelling the role of epigenetic modifications in development and reproduction of angiosperms: a critical appraisal. Front Genet 13:819941

Lalge AB, Mendel P, Vyhnanek T, et al (2016) Effects of different morphoregulators on growth and development of Cannabis sativa L. In: Proceedings of the International Ph. D. Students Conference on MendelNet. pp 726–730

Lapierre É, Monthony AS, Torkamaneh D (2023) Genomics-based taxonomy to clarify cannabis classification. Genome 66:202–211. 10.1139/gen-2023-0005

Li L, Deng X, Zhang T, et al (2022) Propagation methods decide root architecture of Chinese fir: Evidence from tissue culturing, rooted cutting and seed germination. Plants 11:2472

Lisson SN, Mendham NJ, Carberry PS (2000) Development of a hemp (Cannabis sativa L.) simulation model 2. The flowering response of two hemp cultivars to photoperiod. Aust J Exp Agric 40:413–417

Loiseau V, Herniou EA, Moreau Y, et al (2020) Wide spectrum and high frequency of genomic structural variation, including transposable elements, in large double-stranded DNA viruses. Virus Evol 6:vez060

Luo X, Yin M, He Y (2021) Molecular genetic understanding of photoperiodic regulation of flowering time in Arabidopsis and soybean. Int J Mol Sci 23:466

McKernan KJ, Helbert Y, Kane LT, et al (2020) Sequence and annotation of 42 cannabis genomes reveals extensive copy number variation in cannabinoid synthesis and pathogen resistance genes. BioRxiv 2020–2021

McPartland JM (2018) Cannabis systematics at the levels of family, genus, and species. Cannabis Cannabinoid Res 3:203–212

McPartland JM, Guy GW (2017) Models of Cannabis taxonomy, cultural bias, and conflicts between scientific and vernacular names. The botanical review 83:327–381

Mendel P, Schiavo-Capri E, Lalge AB, et al (2020) Evaluation of selected characteristics in industrial hemp after phytohormonal treatment. Pak J Agric Sci 57:1–7

Michael T, Lynch R, Padgitt-Cobb L, et al (2024) Domesticated cannabinoid synthases amid a wild mosaic cannabis pangenome

Michael TP, Salome PA, Yu HJ, et al (2003) Enhanced fitness conferred by naturally occurring variation in the circadian clock. Science (1979) 302:1049–1053

Minitab L (2021) Minitab, LLC. www.minitab.com. Accessed 10 Nov 2023

Montesinos-López OA, Montesinos-López A, Crossa J, et al (2018) Multi-trait, multi-environment deep learning modeling for genomic-enabled prediction of plant traits. G3: Genes, genomes, genetics 8:3829–3840

Monthony AS, de Ronne M, Torkamaneh D (2024) Exploring ethylene-related genes in Cannabis sativa: implications for sexual plasticity. Plant Reprod 37:321–339

Morales F, Ancín M, Fakhet D, et al (2020) Photosynthetic metabolism under stressful growth conditions as a bases for crop breeding and yield improvement. Plants 9:88

Mora-Poblete F, Maldonado C, Henrique L, et al (2023) Multi-trait and multi-environment genomic prediction for flowering traits in maize: a deep learning approach. Front Plant Sci 14:1153040

Naik YD, Bahuguna RN, Garcia Caparros P, et al (2025) Exploring the multifaceted dynamics of flowering time regulation in field crops: Insight and intervention approaches. Plant Genome 18:e70017

Naim-Feil E, Elkins AC, Malmberg MM, et al (2023) The Cannabis Plant as a Complex System: Interrelationships between Cannabinoid Compositions, Morphological, Physiological and Phenological Traits. Plants 12:493

Naim-Feil E, Pembleton LW, Spooner LE, et al (2021) The characterization of key physiological traits of medicinal cannabis (Cannabis sativa L.) as a tool for precision breeding. BMC Plant Biol 21:294

Najafabadi MY, Heidari A, Rajcan I (2023) AllInOne Pre-processing: A comprehensive preprocessing framework in plant field phenotyping. SoftwareX 23:101464

Obeso JR (2002) The costs of reproduction in plants. New phytologist 155:321–348

Osnato M, Cota I, Nebhnani P, et al (2022) Photoperiod control of plant growth: flowering time genes beyond flowering. Front Plant Sci 12:805635

Pantin F, Simonneau T, Muller B (2012) Coming of leaf age: control of growth by hydraulics and metabolics during leaf ontogeny. New Phytologist 196:349–366

Petit J, Salentijn EMJ, Paulo M-J, et al (2020a) Genetic architecture of flowering time and sex determination in hemp (Cannabis sativa L.): a genome-wide association study. Front Plant Sci 11:569958

Petit J, Salentijn EMJ, Paulo M-J, et al (2020b) Genetic variability of morphological, flowering, and biomass quality traits in hemp (Cannabis sativa L.). Front Plant Sci 11:102

Prasad AM, Iverson LR, Liaw A (2006) Newer classification and regression tree techniques: bagging and random forests for ecological prediction. Ecosystems 9:181–199

Purcell S, Neale B, Todd-Brown K, et al (2007) PLINK: a tool set for whole-genome association and population-based linkage analyses. The American journal of human genetics 81:559– 575

Romagnoni A, Jégou S, Van Steen K, et al (2019) Comparative performances of machine learning methods for classifying Crohn Disease patients using genome-wide genotyping data. Sci Rep 9:10351

Salentijn EMJ, Petit J, Trindade LM (2019) The complex interactions between flowering behavior and fiber quality in hemp. Front Plant Sci 10:614

Salentijn EMJ, Zhang Q, Amaducci S, et al (2015) New developments in fiber hemp (Cannabis sativa L.) breeding. Ind Crops Prod 68:32–41

San Nicolas M, Villate A, Alvarez-Mora I, et al (2024) NIR-hyperspectral imaging and machine learning for non-invasive chemotype classification in Cannabis sativa L. Comput Electron Agric 217:108551

Sawler J, Stout JM, Gardner KM, et al (2015) The genetic structure of marijuana and hemp. PLoS One 10:e0133292

Schilling S, Dowling CA, Shi J, et al (2020) The cream of the crop: Biology, breeding and applications of Cannabis sativa. Authorea Preprints

Schwabe AL, McGlaughlin ME (2019) Genetic tools weed out misconceptions of strain reliability in Cannabis sativa: implications for a budding industry. J Cannabis Res 1:1–16

Senan EM, Abunadi I, Jadhav ME, Fati SM (2021) Score and Correlation Coefficient Based Feature Selection for Predicting Heart Failure Diagnosis by Using Machine Learning Algorithms. Comput Math Methods Med 2021:8500314

Shephard H, Parker J, Darby P, Charles CA (2004) Sex expression in hop (Humulus lupulus L. and H. japonicus Sieb. et Zucc.) floral morphology and sex chromosomes. In: Sex determination in plants. Garland Science, pp 139–150

Shi M, Wang C, Wang P, et al (2023) Role of methylation in vernalization and photoperiod pathway: a potential flowering regulator? Hortic Res 10:uhad174

Shi M, Wang C, Wang P, et al (2022a) Methylation in DNA, histone, and RNA during flowering under stress condition: A review. Plant Science 324:111431

Shi Y, Guo E, Cheng X, et al (2022b) Effects of chilling at different growth stages on rice photosynthesis, plant growth, and yield. Environ Exp Bot 203:105045

Small E, Beckstead HD (1973) Cannabinoid phenotypes in Cannabis sativa. Nature 245:147–148

Song YH, Shim JS, Kinmonth-Schultz HA, Imaizumi T (2015) Photoperiodic flowering: time measurement mechanisms in leaves. Annu Rev Plant Biol 66:441–464

Spielmann M, Lupiáñez DG, Mundlos S (2018) Structural variation in the 3D genome. Nat Rev Genet 19:453–467

Stack GM, Toth JA, Carlson CH, et al (2021) Season long characterization of high cannabinoid hemp (Cannabis sativa L.) reveals variation in cannabinoid accumulation, flowering time, and disease resistance. GCB bioenergy 13:546–561

Statista (2024) Cannabis—Worldwide|Statista Market Forecast. https://www.statista.com/outlook/hmo/cannabis/worldwide. Accessed 22 Nov 2024

Steed G, Ramirez DC, Hannah MA, Webb AAR (2021) Chronoculture, harnessing the circadian clock to improve crop yield and sustainability. Science (1979) 372:eabc9141

Steel L, Welling M, Ristevski N, et al (2023) Comparative genomics of flowering behavior in Cannabis sativa. Front Plant Sci 14:1227898

Sultan SE (2000) Phenotypic plasticity for plant development, function and life history. Trends Plant Sci 5:537–542

Tan F, Swain SM (2006) Genetics of flower initiation and development in annual and perennial plants. Physiol Plant 128:8–17

Tang Z, Wang X, Xiang Y, et al (2024) Application of hyperspectral technology for leaf function monitoring and nitrogen nutrient diagnosis in soybean (Glycine max L.) production systems on the Loess Plateau of China. European Journal of Agronomy 154:127098

Team RC (2020) RA language and environment for statistical computing, R Foundation for Statistical. Computing

Torkamaneh D, Laroche J, Belzile F (2020) Fast-GBS v2. 0: an analysis toolkit for genotyping-by-sequencing data. Genome 63:577–581

Torkamaneh D, Laroche J, Boyle B, et al (2021) A bumper crop of SNPs in soybean through high density genotyping by sequencing (HD GBS). Plant Biotechnol J 19:860

Toth JA, Stack GM, Carlson CH, Smart LB (2022) Identification and mapping of major-effect flowering time loci Autoflower1 and Early1 in Cannabis sativa L. Front Plant Sci 13:991680

Turck F, Fornara F, Coupland G (2008) Regulation and identity of florigen: FLOWERING LOCUS T moves center stage. Annu Rev Plant Biol 59:573–594

Warf B (2014) High points: an historical geography of cannabis. Geogr Rev 104:414–438

Webb AAR (2003) The physiology of circadian rhythms in plants. New Phytologist 160:281–303

Webb AAR, Seki M, Satake A, Caldana C (2019) Continuous dynamic adjustment of the plant circadian oscillator. Nat Commun 10:550

Wei H, Chen S, Niyitanga S, et al (2022) Genome-wide identification and expression analysis response to GA3 stresses of WRKY gene family in seed hemp (Cannabis sativa L). Gene 822:146290

Wei T, Simko V, Levy M, et al (2017) Package ‘corrplot.’ Statistician 56:e24

Wu G, Poethig RS (2006) Temporal regulation of shoot development in Arabidopsis thaliana by miR156 and its target SPL3

Zhang M, Anderson SL, Brym ZT, Pearson BJ (2021) Photoperiodic flowering response of essential oil, grain, and fiber hemp (Cannabis sativa L.) cultivars. Front Plant Sci 12:694153

