## Supplemental Figures S1 to S16 for "Predicting flowering time using integrated morphophysiological and genomic data with machine learning models"

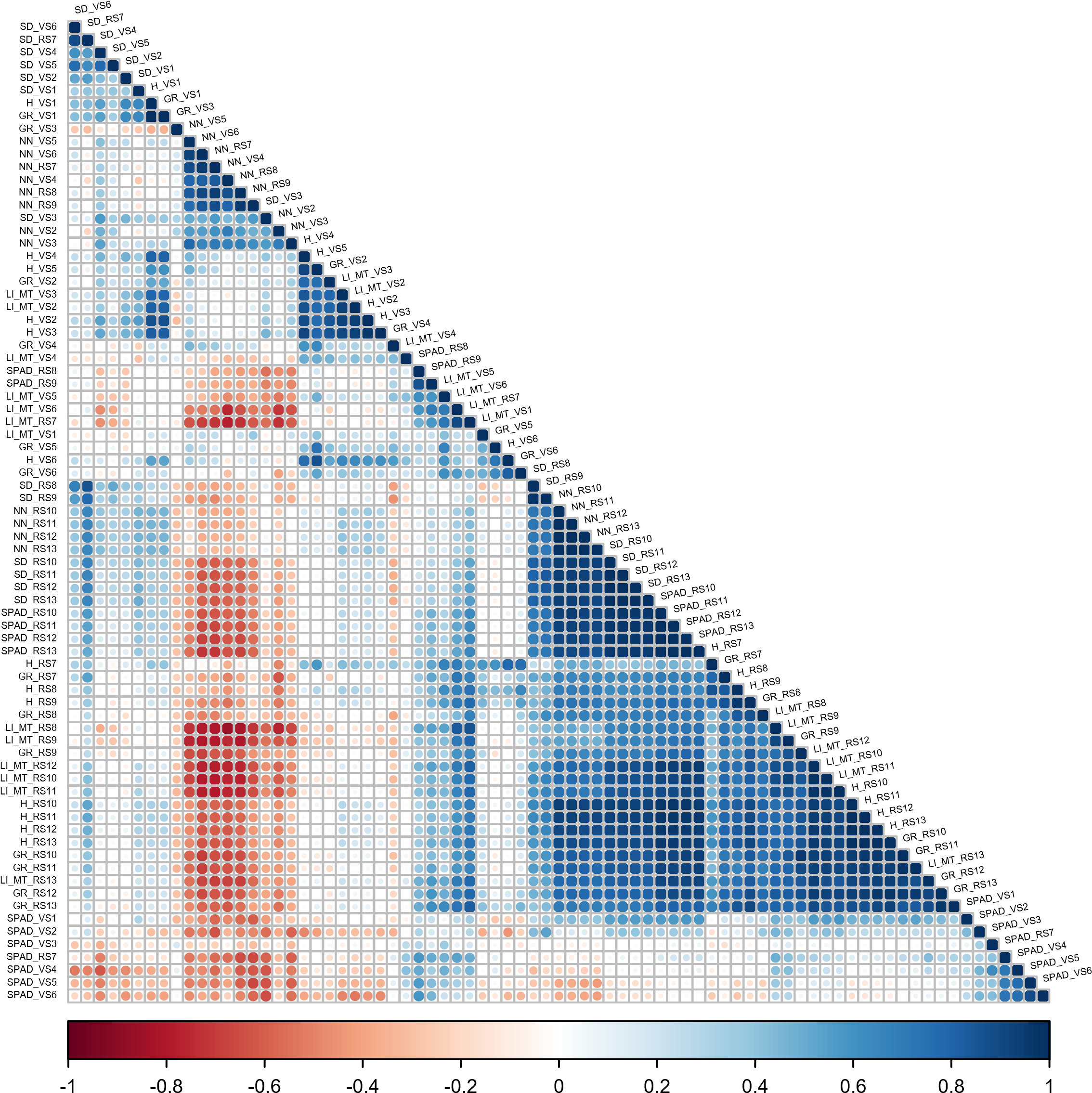

**Fig S1.** Trait correlation heatmap between six key morphophysiological traits in 25 native cannabis populations in Iran, assessed weekly during both VS and RS (13 weeks) based on females. Abbreviations; SD: Stem Diameter, H: Height, GR: Growth Rate, NN: Number of Nodes on the Main Stem, LI-MT: Length of Internode in the Middle Third of the Main Stem, SPAD: SPAD-based Chlorophyll, VS: Vegetative Stage, RS: Reproductive Stages, and the numbers alongside the trait abbreviations indicate the week of data collection.

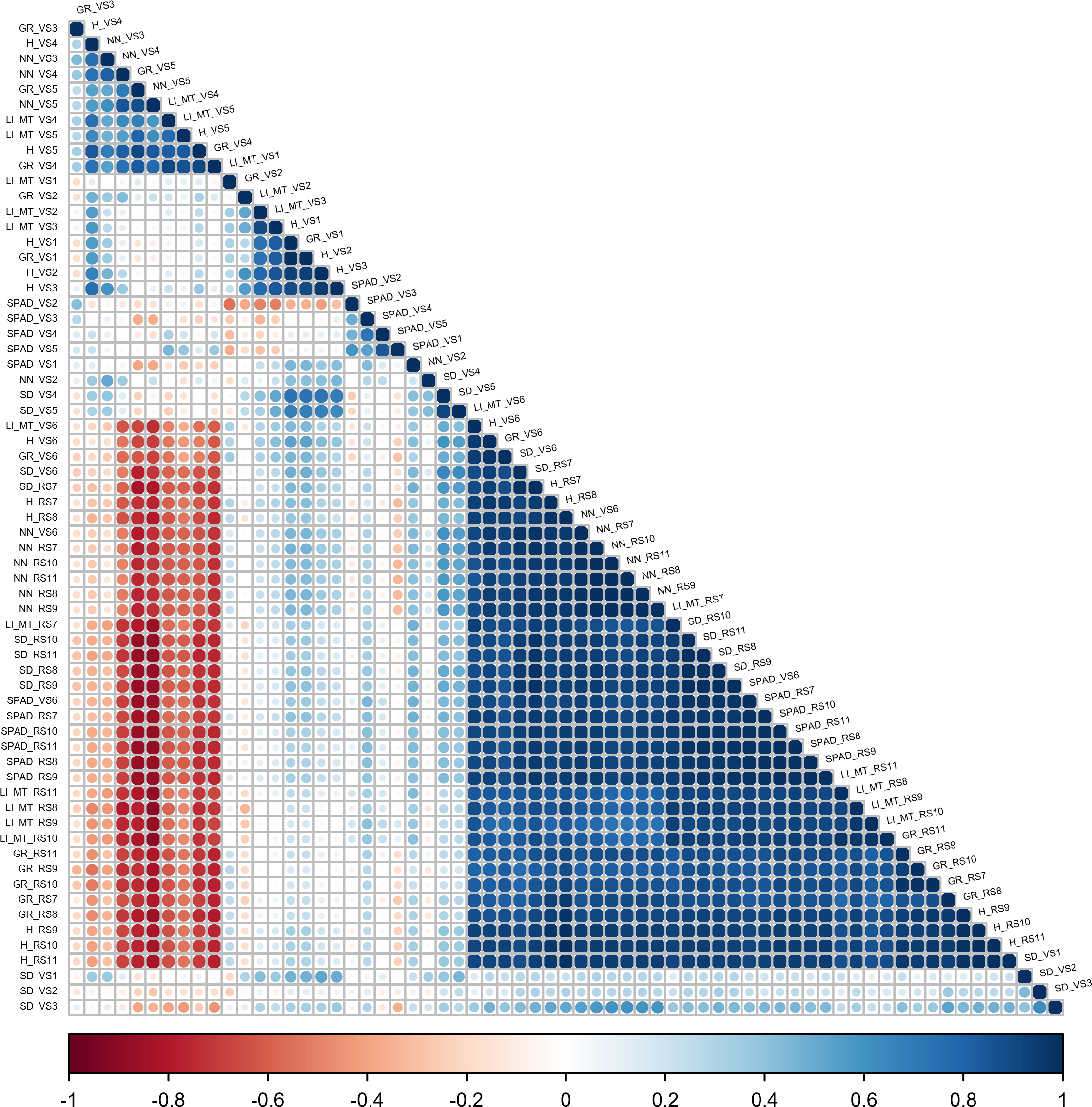

**Fig S2.** Trait correlation heatmap between six key morphophysiological traits in 25 native cannabis populations in Iran, assessed weekly during both VS and RS (11 weeks) based on males. Abbreviations; SD: Stem Diameter, H: Height, GR: Growth Rate, NN: Number of Nodes on the Main Stem, LI-MT: Length of Internode in the Middle Third of the Main Stem, SPAD: SPAD-based Chlorophyll, VS: Vegetative Stage, RS: Reproductive Stages, and the numbers alongside the trait abbreviations indicate the week of data collection.

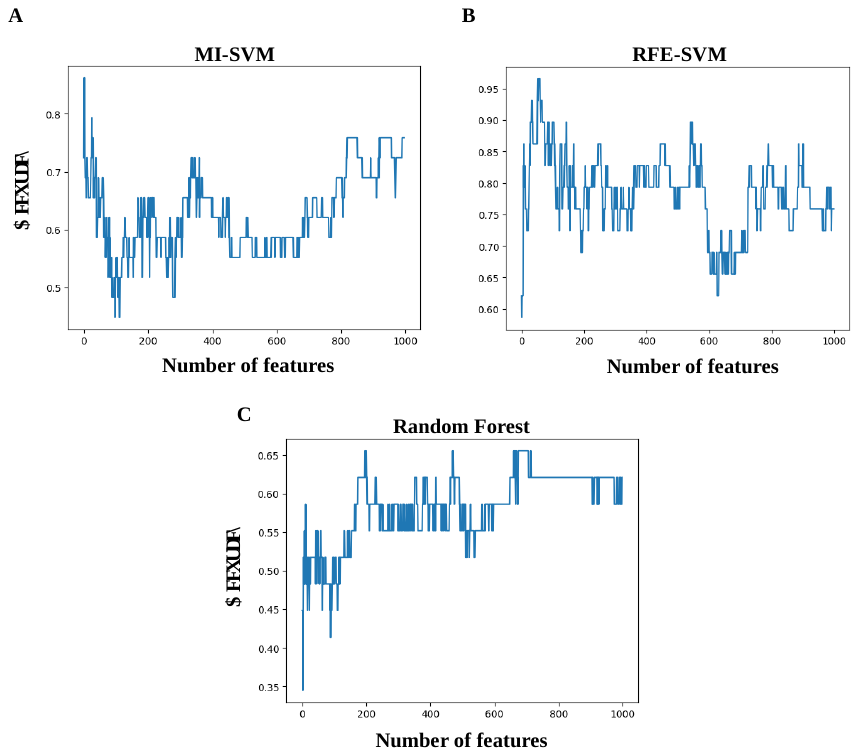

**Fig S3** Feature selection (FS) curves for MI-SVM, RFE-SVM, and RF models are depicted. The number of features, along with the corresponding SVM model weights (**A** and **B**), is illustrated, while the number of trees versus accuracy for the RF model is shown (**C**). (**A**) The peak of the curve is achieved with four features, resulting in 86.2% accuracy. (**B**) The peak of the curve is attained with 53 features, yielding an accuracy of 96.6%. (**C**) Using 197 trees, RF model identifies 2840 features, achieving an accuracy of 65.5%.

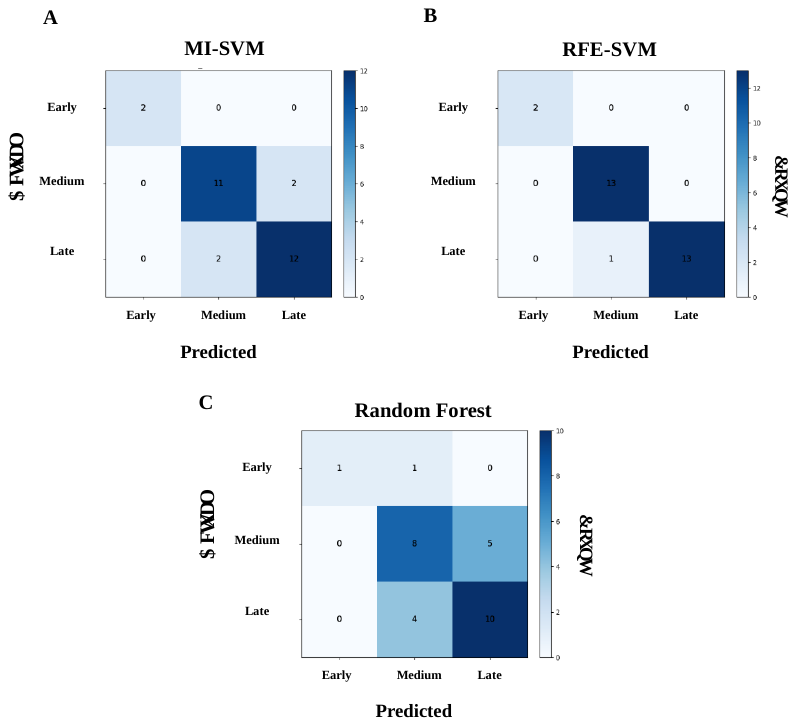

**Fig S4.** Confusion matrix heatmap for different feature selection techniques. Illustrating the distribution of instances across different classes. (**A**) MI-SVM, (**B**) RFE-SVM, and (**C**) RF. The matrix is based on the test set, representing 20% of the total data.

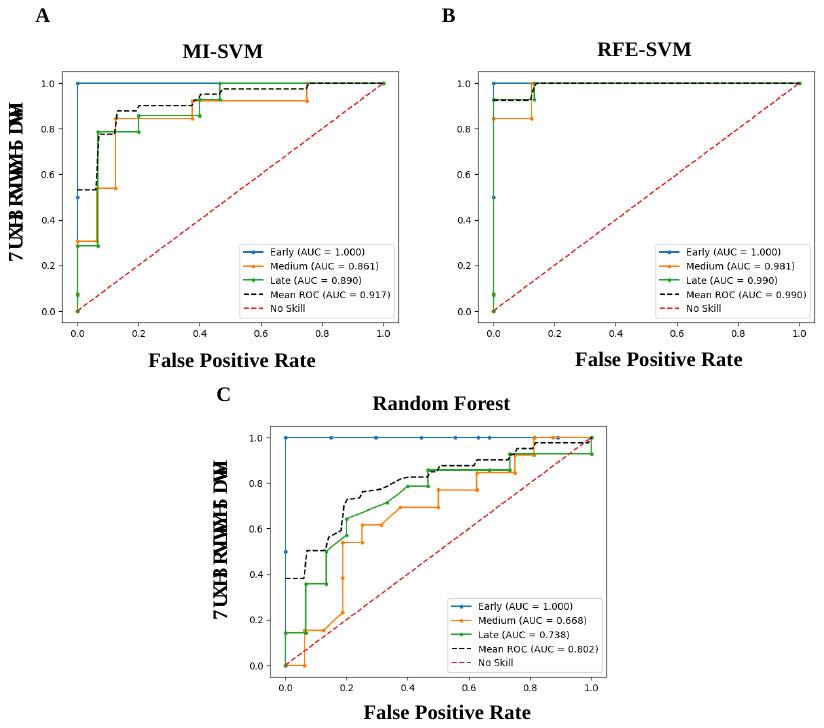

**Fig S5.** ROC AUC curves for different feature selection techniques. (**A**) MI-SVM, (**B**) RFE-SVM, and (**C**) RF. ROC AUC curves are based on the test set, representing 20% of the total data.

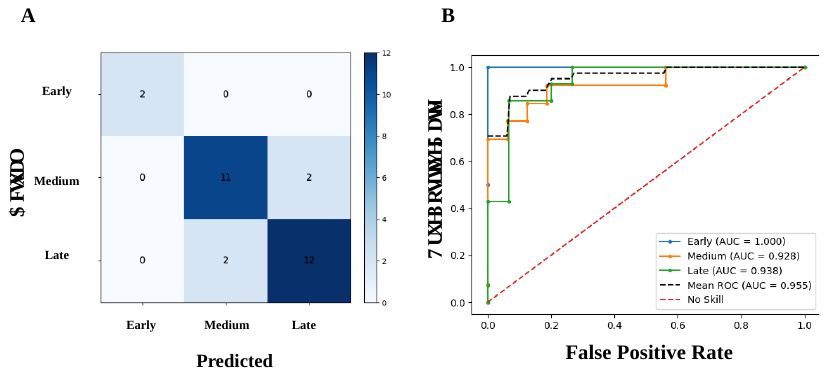

**Fig S6.** (**A**) Confusion matrix heatmap and (**B**) ROC AUC validation using the proposed model with 28 features common between at least two selection techniques. ROC AUC curve and confusion matrix are based on the test set, representing 20% of the total data.

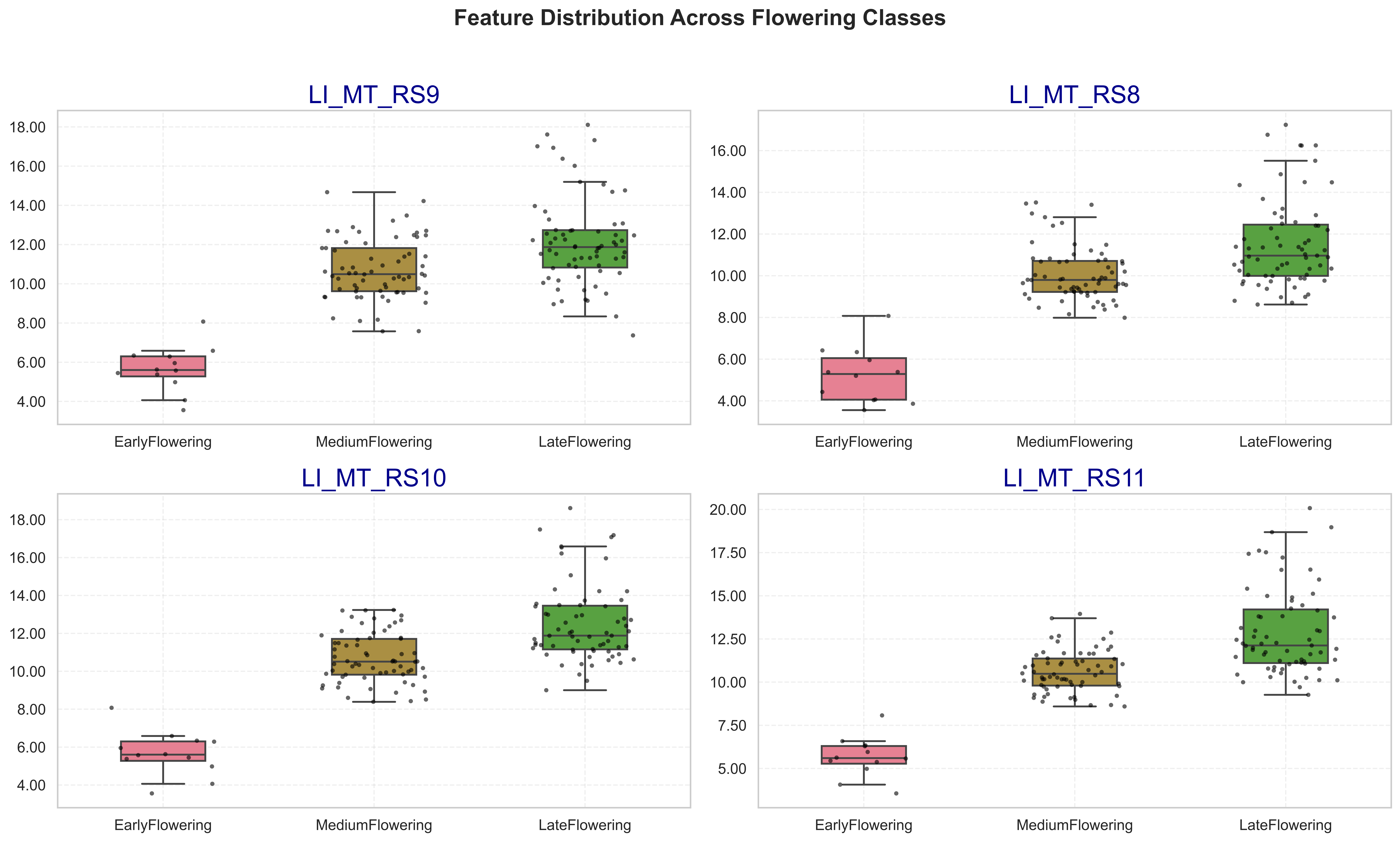

**Fig S7.** Distribution of the four core features (feature set a) across early, medium, and late flowering classes. These features were consistently selected by all three feature selection methods (MI-SVM, RFE-SVM, and RF) and enabled 86.2% classification accuracy with high class separation.

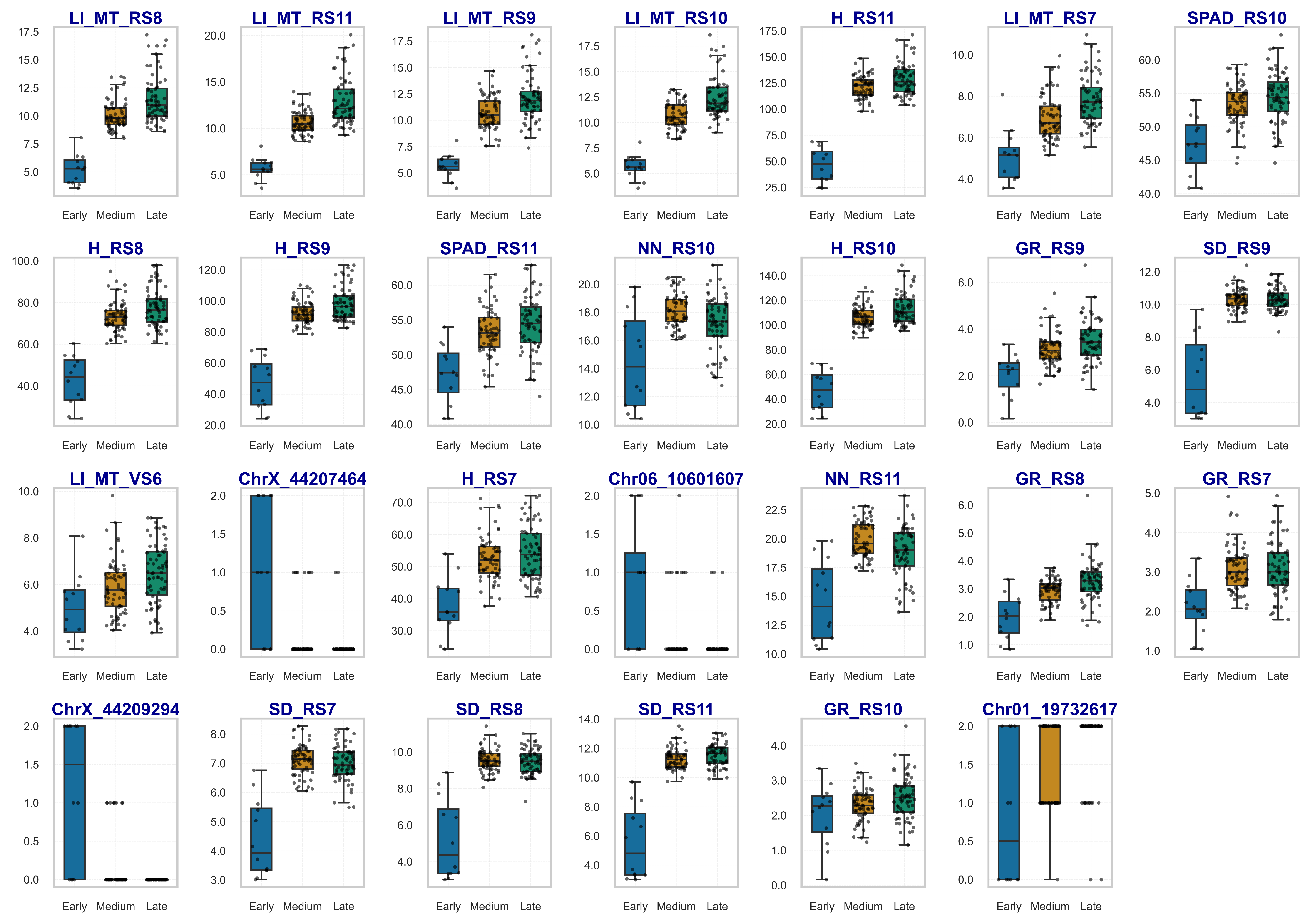

**Fig S8.** Boxplots of the 28 features are selected by at least two methods (feature set b), including the four shared features. This intermediate set maintained 86.2% accuracy but showed improved ROC AUC (0.955), capturing more nuanced phenotypic variation.

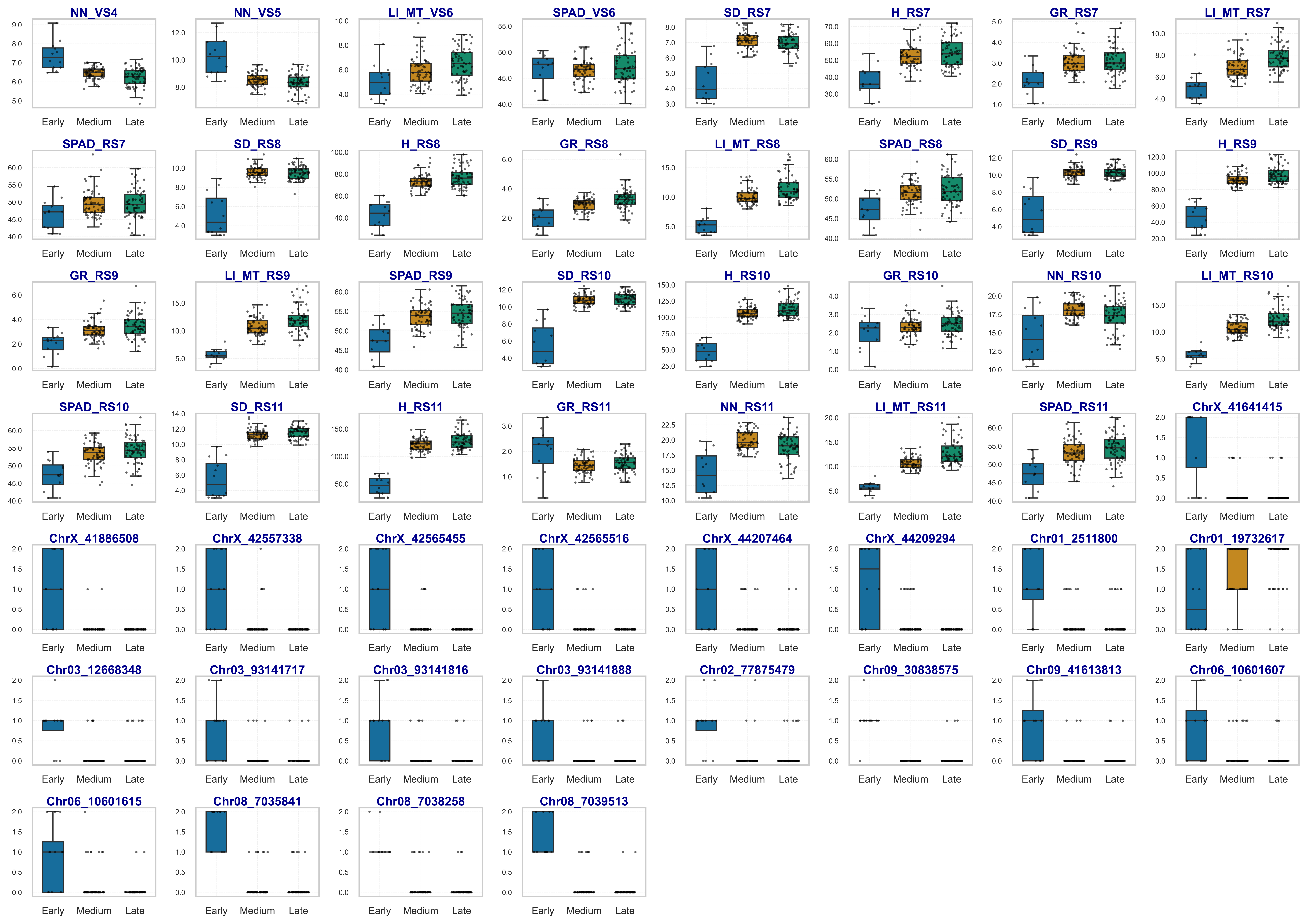

**Fig S9.** Distribution of the 53 features selected exclusively by RFE-SVM (feature set c), combining dynamic morphophysiological traits and genome-wide SNPs. This comprehensive set achieved the highest performance (96.6% accuracy, AUC = 0.994) and highlights the power of integrating phenotypic and genetic information.

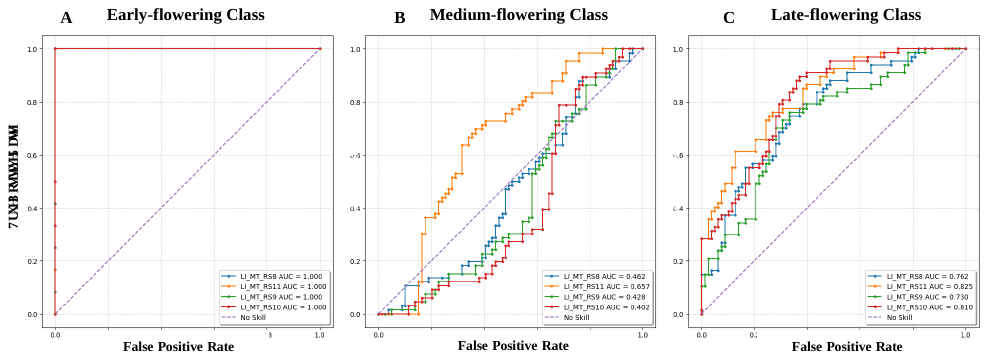

**Fig S10.** ROC AUC analysis showing the discriminative potential of markers identified from features common between all selection techniques (feature = 4, accuracy = 91.7%). This analysis presents the ROC curves for the early-flowering (**A**), medium-flowering (**B**), and late-flowering (**C**) classes. Higher AUC values indicate a higher ability of the identified markers to distinguish between the classes. The analysis was conducted based on 5-fold cross-validation in the proposed model.

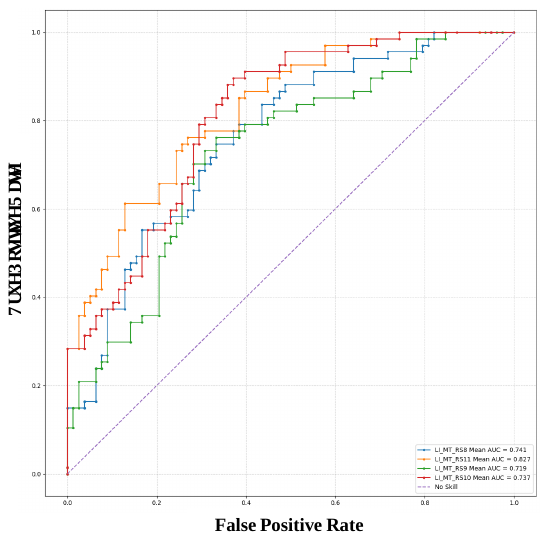

**Fig S11.** ROC AUC curve represents the average discriminative potential of markers identified from features common between all selection techniques (feature = 4, accuracy = 91.7%) the early-flowering, medium-flowering, and late-flowering classes. The x-axis shows the false positive rate (FPR), and the y-axis indicates the true positive rate (TPR). The average AUC value reflects the overall ability of the identified markers to distinguish between the classes. This analysis was conducted using 5-fold cross-validation in the proposed model.

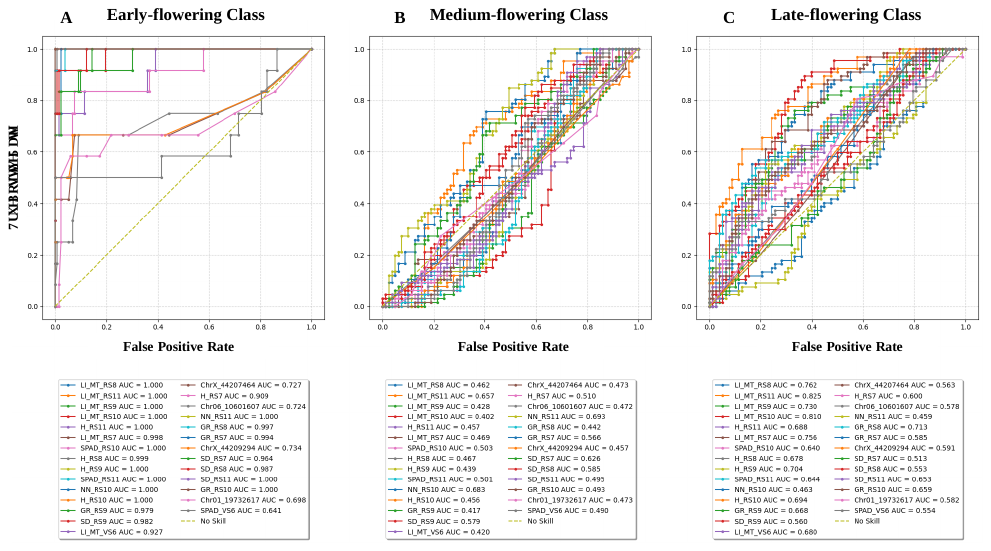

**Fig S12.** ROC AUC analysis showing the discriminative potential of markers identified from features common between at least two selection techniques (feature = 28, accuracy = 95.5%). This analysis presents the ROC curves for the early-flowering (**A**), medium-flowering (**B**), and late-flowering (**C**) classes. Higher AUC values indicate a higher ability of the identified markers to distinguish between the classes. The analysis was conducted based on 5-fold cross-validation in the proposed model.

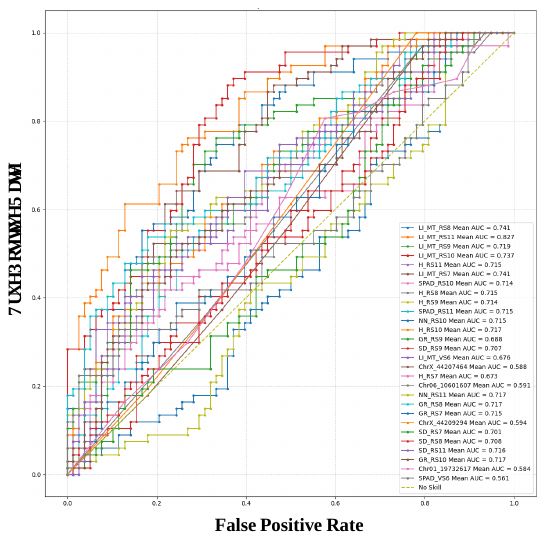

**Fig S13.** ROC AUC curve represents the average discriminative potential of markers identified from features common between at least two selection techniques (feature = 28, accuracy = 95.5%) the early-flowering, medium-flowering, and late-flowering classes. The x-axis shows the false positive rate (FPR), and the y-axis indicates the true positive rate (TPR). The average AUC value reflects the overall ability of the identified markers to distinguish between the classes. This analysis was conducted using 5-fold cross-validation in the proposed model.

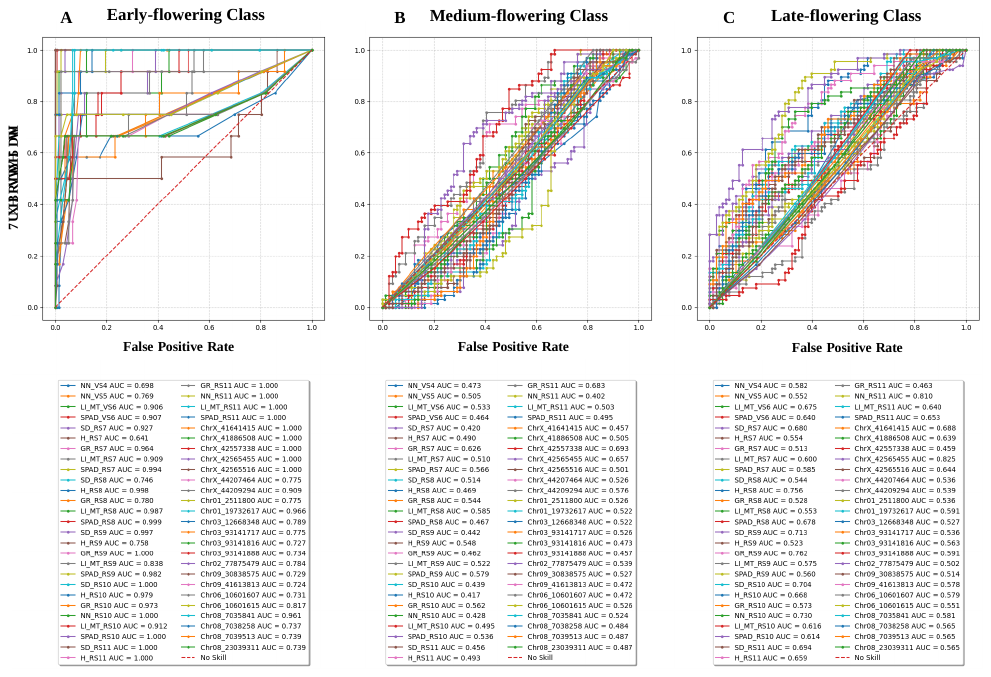

**Fig S14.** ROC AUC analysis showing the discriminative potential of markers identified by the RFE-SVM (feature = 53, ROC AUC = 99.4%). This analysis presents the ROC curves for the early-flowering (**A**), medium-flowering (**B**), and late-flowering (**C**) classes. Higher AUC values indicate a higher ability of the identified markers to distinguish between the classes. The analysis was conducted based on 5-fold cross-validation in the proposed model

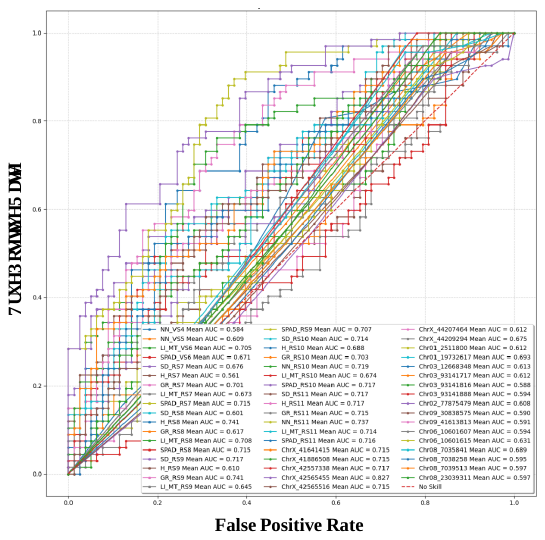

**Fig S15.** ROC AUC curve represents the average discriminative potential of markers identified by the RFE-SVM (feature = 53, ROC AUC = 99.4%) across the early-flowering, medium-flowering, and late-flowering classes. The x-axis shows the false positive rate (FPR), and the y-axis indicates the true positive rate (TPR). The average AUC value reflects the overall ability of the identified markers to distinguish between the classes. This analysis was conducted using 5-fold cross-validation in the proposed model.

**Fig S16.** Haplotype blocks (HB) and linkage disequilibrium (LD) structure among 22 candidate SNP markers associated with flowering time differentiation in cannabis. The plot illustrates both LD values (r² and D') and the organization of haplotype blocks. Color intensity represents LD strength (red = high, white/blue = low). The haploblocks delineate regions of high correlation between SNPs, highlighting potential genomic regions contributing to the genetic differentiation of early, medium, and late flowering classes.
